# Phenotypic and *omics* analyses of the Sordariomycetes *Marquandomyces marquandii* and *Albophoma yamanashiensis* isolated from estuarine sediments

**DOI:** 10.1101/2023.07.26.550687

**Authors:** Z. Agirrezabala, A. Otamendi, R. Liébana, M. Azkargorta, C. Perez-Cruz, M. T. Dueñas, M. Ostra, R. López, I.V. Grigoriev, L. Alonso-Sáez, F. Elortza, A. Lanzén, S. Haridas, Oier Etxebeste

**Affiliations:** Laboratory of Microbiology, Dept. of Applied Chemistry, Faculty of Chemistry, University of the Basque Country (UPV/EHU), 20018 San Sebastian; AZTI, Marine Research, Basque Research and Technology Alliance (BRTA), Sukarrieta; Proteomics Platform, Center for Cooperative Research in Biosciences (CIC bioGUNE), Basque Research and Technology Alliance (BRTA), CIBERehd, 48160, Derio; Laboratory of Analytical Chemistry, Dept. of Applied Chemistry, Faculty of Chemistry, University of the Basque Country (UPV/EHU), 20018 San Sebastian; Dept. of Organic Chemistry I, Faculty of Chemistry, University of the Basque Country (UPV/EHU), 20018 San Sebastian; U.S. Department of Energy Joint Genome Institute (JGI), Lawrence Berkeley National Laboratory, Berkeley, CA 94720, USA; Department of Plant and Microbial Biology, University of California Berkeley, Berkeley, CA 94720, USA; AZTI, Marine Research, Basque Research and Technology Alliance (BRTA), Pasaia; IKERBASQUE, Basque Foundation for Science, Bilbao

**Author notes:** Corresponding author; Lab. of Biology, Dept. of Applied Chemistry, Faculty of Chemistry, University of the Basque Country, Manuel de Lardizabal, 3, 20018, San Sebastian.

**Keywords:** . Estuarine filamentous fungi, Sordariomycetes, Hypocreales, CAZYmes, secondary metabolite gene clusters, secondary metabolites, fucoidan

## Abstract

Marine environments harbor a vast diversity of microorganisms, which have developed multiple strategies to adapt to challenging conditions and represent a valuable source for new products such as pigments, enzymes and bioactive compounds. From all microorganisms inhabiting marine environments, fungi have been the least studied, despite their ubiquitous presence and great biotechnological potential. Here, we focused on the isolation and characterization of filamentous fungi from marine sediment samples, which were collected along the Basque coast in Spain. Through phenotypic characterization, we identified isolates potentially able to produce secondary metabolites or grow on minimal culture medium supplemented with recalcitrant algal polysaccharides. Based on this screening, two Sordariomycetes isolates were selected for further analyses through genome sequencing and *omics* techniques: 1) a *Marquandomyces marquandii* strain able to stain the culture medium in yellow, indicative of secretion of pigments and secondary metabolites and 2) an *Albophoma yamanashiensis* strain able to grow in minimal culture medium supplemented with the recalcitrant algal polysaccharide fucoidan. Fungal co-culture experiments suggested an inhibitory effect of the secretome of *M. marquandii* on fungal growth. Under culture conditions inducing pigment secretion, a set of secondary metabolite gene clusters were differentially expressed, as analysed by RNA-seq. On the other hand, transcriptomic and proteomic experiments on *A. yamanashiensis* unveiled the enzymatic activities expressed in response to the presence of fucoidan. Overall, our results indicate that the isolated marine fungal strains could serve as a source of new enzymatic activities and secondary metabolites.

## 1. Introduction

Fungi probably represent the largest and most diverse eukaryotic kingdom, with an estimated number of species ranging from 2 to 3 million [1–4]. They have adapted to very diverse environments and niches [5], including high salt concentrations, nutrient limitation, or environments where the main source of carbon and nitrogen is in form of recalcitrant polymers. The first marine fungal strains were isolated more than a century ago [6] and research interest on marine mycology has grown considerably in the last years. However, the state-of-the-art in this field lags behind that of other microorganisms such as marine bacteria. As a reference, the number of total marine fungal species listed in the Marine Fungi website (https://www.marinefungi.org/; [7]) is 2,149, as of February 2025. In contrast, the bacterial database BacDive [8] includes a subset of 7,916 strains with the tag “*aquatic*”, 2,040 strains with the tag “*marine*” and 1,550 strains with the tag “*sediment*” (accessed in October 2024). Based on metabarcoding analyses, it is hypothesized that the number of marine fungal species and their diversity is far larger than the number of species currently deposited in culture collections [6,9,10]. One of the reasons explaining this gap is the difficulty for cultivating slow-growing marine fungi under laboratory conditions [11]. Overall, it can be concluded that the potential impact of fungi at environmental and biotechnological levels is underestimated [6,12].

Genome sequencing of cultivable marine fungal strains has revealed biotechnological potential in several fields, such as the identification of compounds with antitumoral activity as well as new antimicrobials, pigments and other secondary metabolites [13]. Fungal secondary metabolites are compounds of different types, such as polyketides, terpenes, non-ribosomal peptides (NRPs), ribosomally synthesized and post-translationally modified peptides or RiPPs, which confer advantages in particular niches [14]. Their synthesis, modification, cellular compartmentalization, and secretion are encoded in the genome by groups of genes that are typically located in clusters. Elucidating the genetic mechanisms controlling the expression of these secondary metabolite gene clusters and the chemical reactions catalyzed by their activities is key for the identification of new bioactive compounds and the optimization of their industrial synthesis.

Marine fungal strains capable of degrading polysaccharides of marine origin have also been identified [13,15,16], which are of interest in biotechnology. Marine polysaccharides, synthesized by phytoplankton and macroalgae during photosynthesis, exhibit a great diversity of chemical structures, in terms of monosaccharide composition, linkage types and decoration patterns [17]. Among these, fucoidans are complex algal polysaccharides mainly composed of fucose, typically branched and heavily sulfated, which confers them negative charges. The degradation of these complex molecules involves a coordinated action of Carbohydrate-Active enZymes (CAZymes) and sulfatases [18]. CAZymes, including glycoside hydrolases (GHs), carbohydrate esterases (CEs), and polysaccharide lyases (PLs) break down fucoidan by hydrolyzing glycosidic bonds, while sulfatases remove sulfate groups to facilitate further degradation. Notably, the knowledge on the ability of marine fungi to degrade fucoidan and the molecular mechanisms employed is still very limited. Discovering new enzymes with novel catalytic activities modifying and breaking down fucoidan is of great interest, since these polysaccharides and the oligosaccharides resulting from their partial degradation have shown immunomodulatory, anti-inflammatory, virucidal or anticancer activities [19–23].

In this work, we focused on the isolation of filamentous fungi from sediment samples collected in five estuaries of the Basque Country (Bay of Biscay). From the resulting library of isolates, mainly formed by Sordariomycetes and Eurotiomycetes fungi, we focused our analysis on isolate M60 (*Marquandomyces marquandii*) based on its ability to produce and secrete pigments, which may be used in the food, cosmetic and textile sectors [24]; and M98 (*Albophoma yamanshiensis*), due to its phenotype on culture medium supplemented with polysaccharides of marine origin. Results of phenotypic and *omics* analyses of both isolates strongly suggest that the isolates of our library could serve as a source of new enzymatic activities and secondary metabolites.

## 2. Experimental

### 2.1. Sample collection, processing and isolation of filamentous fungi

Seven stations along the Basque coast (Bay of Biscay) were selected for sampling (Figure 1). The selected samples covered diverse values of organic content and granulometry properties. One of these stations (LUR20) is located approximately 0.8 km from the coast, at a depth of 37 m (black square in Figure 1). Samples were collected during winter and spring of 2021, in 50 mL polypropylene tubes, which were stored at 4 °C until processing.

**Figure 1:**
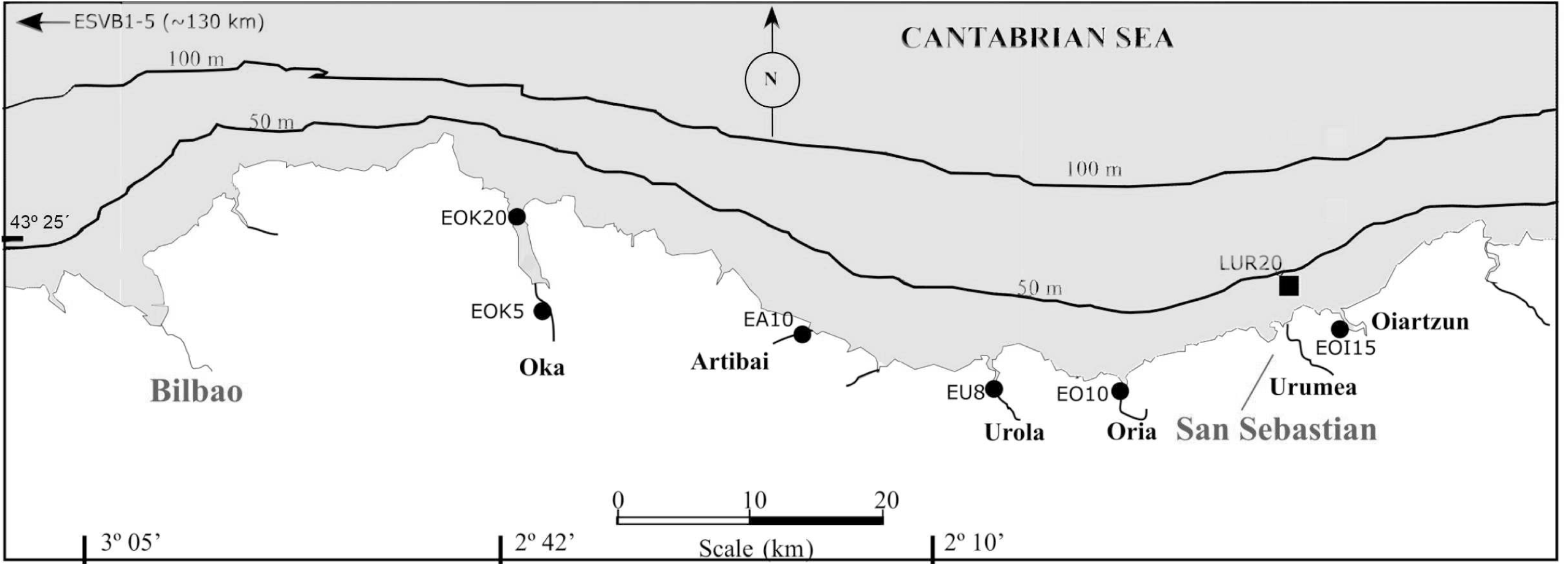
Geographic location of the seven sampling points along the Basque coast selected in this work. Circles indicate estuaries while the square indicates a coastal area. EOK: river Oka; EA: river Artibai; EU: river Urola; EO: river Oria; LUR: river Urumea; EOI: river Oiartzun.

For isolation of fungi, the *dilution-to-extinction* method was used [11] based on the procedure developed by Velez and coworkers [25]. Briefly, approximately two centimeters of the upper layer of the sediment were removed using a sterile spatula. Samples of 1.4-2.2 g were taken from the center of each tube (material that was not in contact with the wall of the tube) and transferred to 15 mL polypropylene tubes. Sediment samples were diluted serially (1/5 once and 1/10 twice) in artificial sea water (ASW; see below). Aliquots of 50 µL per well were inoculated in 24-well plates previously filled with the culture medium (see below) plus penicillin. One well per plate was inoculated with ASW as a control for potential contaminations. Plates were covered with aluminum foil and cultured under aerobic conditions at atmospheric pressure and 25° C. Growth was checked regularly and samples of isolates generating filaments were transferred to plates containing the same culture medium used for isolation, until homogeneous phenotypes, and thus probably axenic cultures, were obtained. Specific isolates showed invasive behavior and colonized adjacent wells in multi-well plates. To minimize contamination, these isolates were discarded from the analysis. Samples of all selected isolates were stored at -80° C in a glycerol solution (18% in artificial seawater).

### 2.2. Preparation of culture media and solutions

ASW solutions were prepared by diluting 90 g of sea salts (Millipore) per liter of distilled water (3x stock; 9%; ASW1) [15]. Alternatively, a 4x stock (ASW2) was prepared by adding 46.94 g NaCl, 7.84 g Na_2_SO_4_, 21.28 g MgCl_2_·6H_2_O, 2.20 g CaCl_2_, 0.38 g NaHCO_3_, 1.33 g KCl, 0.19 g KBr, 0.05 g H_3_BO_3_, 0.05 g SrCl_2_ and 0.006 g NaF per liter of distilled water [26]. Penicillin sodium salt (0.018 mg/mL; 30 units/mL) was added to all culture media used for isolation. Six types of media carrying different carbon and nitrogen sources were used for isolation of filamentous microbes. In a first screen, the following culture media (1/5 dilution of the standard; 15 g of agar per liter) were used: 1) malt extract agar (MEA; Condalab; 2.69 g of the powder, 133 mL of ASW1 or 100 mL of ASW2 solutions and distilled water to 400 mL); 2) potato dextrose agar (PDA; Condalab; 3.12 g of the powder, 133 mL of ASW1 or 100 mL of ASW2 solutions and distilled water to 400 mL); 3) corn meal agar (CMA; Condalab; 1.36 g of the powder, 133 mL of ASW1 or 100 mL of ASW2 solutions and distilled water to 400 mL); 4) Lc medium (2g Arabic gum, 2 g olive oil, 0.2 g yeast extract, 0.4 g peptone, 2.42 g PIPES (1,4-Piperazinediethanesulfonic acid), 133 mL of ASW1 or 100 mL of ASW2 solutions and distilled water to 400 mL; pH adjusted to 7.5 with NaOH); and 5) Pc medium (3.6 g brain heart infusion, 9.2 g NaCl, 2.42 g PIPES, 133 mL of ASW1 or 100 mL of ASW2 solutions and distilled water to 400 mL; pH adjusted to 7.5 with NaOH) [25,27,28].

A minimal medium (MMM) supplemented with different carbon and nitrogen sources was also prepared by adding, per liter, 250 mL of ASW2 solution and 10 mL of a salts plus trace elements solution (25 mL of a trace elements solution, 50 g KH_2_PO_4_, 10 g MgSO_4_.7H_2_O, 5 g CaCl_2_·2H_2_O per liter) [15]. The trace elements solution (40x) was prepared by adding, per liter, 20 g monohydrated citric acid, 20 g ZnSO_4_.7H_2_O, 2.84 g FeSO_4_·7H_2_O, 0.15 g MnSO_4_·H_2_O, 1.04 g CuSO_4_·5H_2_O, 0.2 g H_3_BO_3_, 0.15 g (NH_4_)_6_Mo_7_O_24_·4H_2_O and 0.2 g biotin.

### 2.3. Phenotypic screening of the fungal isolates

To ease the screening of isolates, we decided to use solid MMM. First, agar concentration had to be decreased as low as possible because it could be used by the isolates as a carbon source. Agar concentration was decreased from 1.5% [25] to 0.4-0.5%, which was the minimum agar concentration allowing solidification of the culture medium and clean point inoculation (Figure S1). Minimal medium (0.5% agar) without the addition of an extra carbon source was used as a negative control, while the addition of glucose (2 g/L) was used as a positive control of growth. The commercial algal polysaccharides alginate (Sigma-Aldrich; G/M, guluronic acid (G) versus mannuronic acid (M), ratio not provided by the supplier) or fucoidan (*Laminaria japonica*, Carbosynth; *Fucus vesiculosus*, Sigma-Aldrich; or *Undaria pinnatifida*, Sigma-Aldrich) were added at a concentration of 2 g/L (20 g/L stocks). The phenotypes in these culture media were compared to those shown in minimal medium with the β-glucan polysaccharide laminarin (from *Thallus laminariae*; Carbosynth; ratio of β-1,3 and β-1,6 glycosidic linkages not determined by the supplier; also, at 2 g/L), which in general is more easily degraded by microorganisms [13]. Ammonium tartrate (9.2 g/100 mL; 100x) or sodium nitrate (4.25 g/100 mL; 100x) were added as the main nitrogen sources, to compare any possible effect of media acidification (ammonium) or alkalinization (nitrate) in the growth of fungal isolates, and also a hypothetic use of tartrate as a carbon source. Overall, we considered an isolate for further analysis when it showed greater radial growth rate on minimal medium supplemented with the polysaccharide than on minimal medium without an extra carbon source (negative control; only bacteriological agar). This phenotype had to be observed both when ammonium tartrate or sodium nitrate was used as the nitrogen source.

### 2.4. Extraction of genomic DNA, ITS amplification and Sanger sequencing

Isolates were cultured in liquid minimal medium (28° C, 170 rpm) supplemented with ammonium tartrate and glucose as nitrogen and carbon sources. Incubation times varied depending on the growth rate of each isolate. Mycelia were filtered using Miracloth paper and lyophilized. Samples were processed with a Next Advance Bullet Blender homogenizer and genomic DNA fractions were extracted following the 1) phenol:chloroform:isoamylalcohol (25:24:1; Panreac AppliChem) method or, alternatively, 2) the guidelines of the NucleoSpin Plant II Mini kit (Macherey-Nagel), which uses hexadecyltrimethylammonium bromide (CTAB) in the lysis buffer [29]. The integrity of the samples was checked by agarose electrophoresis (0.8%).

Using genomic DNA samples as template, ITS regions or a fragment of the gene *rpb1*, which encodes the main subunit of the RNA polymerase II complex, were amplified using oligonucleotide pairs ITS5 (GGAAGTAAAAGTCGTAACAAGG) and ITS4 (TCCTCCGCTTATTGATATGC), ITS4 and ITS1 (TCCGTAGGTGAACCTGCGG), and/or Rpb1-Up (GAATGCCCAGGTCATTTCGG) and Rpb1-Dw (CCAGCGATATCGTTGTCCATATA) [25,30,31]. The Kapa Taq DNA polymerase (Kapa Biosystems) was used for amplification (denaturation: 30 seconds at 95° C; annealing: 30 seconds at 54° C; elongation: 60 seconds at 72° C; 25 cycles). Amplification and specificity of the amplicons were checked by agarose electrophoresis (2%). Amplicons were purified using the Macherey-Nagel NucleoSpin gel and PCR clean[up kit and Sanger sequenced at Secugen S.L. (Madrid, Spain). The Sankey diagram was generated at SankeyMatic (https://sankeymatic.com/).

### 2.5. Genome sequencing and analysis

Genomes of isolates M60 and M98 were initially sequenced using the Illumina MiSeq platform at the General Services of the University of the Basque Country (UPV/EHU), and later re-sequenced at the CNAG (https://www.cnag.crg.eu/), combining the Illumina NovaSeq platform and Oxford Nanopore Technologies GridION sequencing. Samples were quality-checked at the UPV/EHU and CNAG facilities before sequencing. See data availability for the two whole genome shotgun projects in section 9.

Galaxy (v22.05.1) was used to process genome-sequencing data [32]. The workflow *Itsas_Genomak* can be accessed in the following link: https://usegalaxy.org/u/oilier/h/itsasgenomak. Spades (versions v3.13.1) was used for hybrid genome assembly combining all Illumina and Nanopore sequencing files [33]. The provided assembly was modified to comply with JGI standards by removal of contaminants, organellar sequences, and small unsupported scaffolds, if present. The JGI Fungal Annotation pipeline [34,35] was used to predict genes, provide functional annotations and carry out KOG (EuKaryotic Orthologous Groups) analyses, which were compared with those of closely related species. The trees in the heatmaps showing KOG analyses were generated, using the IDs of the species analyzed, with the common tree function at the NCBI taxonomy browser, and were formatted using iTOL [36]. A mitochondrial scaffold in each dataset was extracted from the main assembly and is presented separately as the mitochondrial assembly. Both datasets are accessible at https://mycocosm.jgi.doe.gov/Marma1 for *M. marquandii* and https://mycocosm.jgi.doe.gov/Albya1 for *A. yamanashiensis* [34].

BUSCO (Benchmarking Universal Single-Copy Orthologs; v5.2.2; hypocreales_odb10 dataset) was used for the assessment of the completeness of genome assemblies [37,38]. ANI (average nucleotide identity) and AAI (average amino acid identity) for species proximity were carried out at enveomics [39] and with AAI-profiler [40] (both websites were accessed in May 2023 and December 2024). The genomes and proteomes used in ANI and AAI analyses were downloaded from the NCBI database (Table S1). CAZyme analyses were carried out at the dbCAN3 website (accessed in December 2024 [41,42]) and results were compared to those provided by MycoCosm, which were obtained after a semi-manual curation of protein filtered model sequences by the CAZy team [43]. The dbCAN3 website uses three tools for automated CAZyme annotation: HMMER for annotated CAZyme domain boundaries, DIAMOND for fast blast hits in the CAZy database, and HMMER for dbCAN-sub, a database of carbohydrate active enzyme subfamilies for substrate annotation. Following the recommendation of the dbCAN3 tool, we excluded from our analyses all the CAZymes predicted only by one of those three tools (1674 and 1403 in the case of M60 and M98, respectively). dbCAN3 also predicted the presence of signal peptides in CAZymes using SignalP. To identify sulfatases, the Sulfatlas website (v2.3.1) was used [44]. The fungal version of AntiSMASH 7.0 was used for prediction of secondary metabolite gene clusters [45].

### 2.6. Phenotypic characterization of isolates M60 and M98

For microscopy analyses, isolates M60 (*Marquandomyces marquandii*) and M98 (*Albophoma yamanashiensis*) were inoculated in liquid MMM supplemented with glucose and ammonium tartrate and cultured at 170 rpm and 28° C. Culture times varied depending on the growth rate of each isolate. Samples of each culture were analyzed using a Nikon Eclipse E100 microscope equipped with an Imaging Source DFK23UP031 camera. Alternatively, a Nikon Eclipse Ci manual upright microscope was used, which included 40× and 100× (oil) Nikon CFI Plan Fluor lenses and a Teledyne Photometrics Moment camera. ImageJ (https://imagej.nih.gov/ij/) (US. National Institutes of Health, Bethesda, MA, USA) was used to process phase contrast images.

For characterization on solid MMM medium, glucose, mannose, fucose or fucoidans from *F. vesiculosus* or *U. pinnatifida* (see above) were used as main carbon sources at a concentration of 2 g/L (20 g/L stocks). Tolerance to salt stress was tested by adding increasing concentrations (0.2, 0.5, 1.0, 2.0 and 3 M) of NaCl to the culture medium. Optimal growth temperatures were determined by testing growth of both isolates at 25, 30, 35 and 37° C. Pigment secretion by *M. marquandii* was assessed and co-cultures of fungal strains were carried out on PDA, MEA, *Aspergillus* minimal (AMM) [46] or Yeast extract, Peptone and Dextrose (YPD) media.

For characterization of pigment production, coloring of media in *M. marquandii* cultures was followed by measuring optical density. Briefly, 10^6^ spores/mL were inoculated in Erlenmeyer flasks filled with 250 mL of potato dextrose broth (PDB), malt extract broth (MEB) or liquid AMM (glucose and ammonium tartrate were used in the latter case as carbon and nitrogen sources, respectively). Three flasks were inoculated per culture medium. Spores were cultured at 25° C. Samples of 2 mL were taken every 24 hours and filtered using Minisart 0,2 µm pore filters (Sartorius). Optical density was measured at 380 nm (NanoDrop 2000c, Thermofisher Scientific), for 14 days.

### 2.6. Co-cultures of fungal isolates

*M. marquandii* was co-cultured with other strains of our collection (M98 – *A. yamanashiensis*; M122 – *genus Acrostalagmus* according to ITS/*rpb1* sequencing data; and MFuc12 – *genus Gliomastix*) as well as with *Aspergillus nidulans* strain TN02A3 [47], on Petri dishes of 14 cm filled with MEA, PDA or AMM culture media. *M. marquandii* was point-inoculated first, and cultured at 25° C. After 96 hours for *A. yamanashiensis* isolate M98 and *Gliomastix* isolate MFuc-12, or 144 hours for *A. nidulans* strain TN02A3 or *Acrostalagmus* isolate M122, the second strain was point-inoculated at a distance of 2.5 cm.

### 2.7. RNA extraction and sequencing

Spores of *A. yamanshiensis* were collected from MEA cultures (168 hours of incubation at 25 °C in the dark) and added (10^6^ spores/mL) to liquid MMM supplemented with ammonium tartrate and one of the following three carbon sources: 1) mannose, as control, 2) fucoidan from *Fucus vesiculosus* or 3) fucoidan from *Undaria pinnatifida*. Spores of *M. marquandii* (10^6^ spores/mL) were also added to MEB and PDB media (malt extract and potato dextrose broth, respectively). The cultures of *M. marquandii* were incubated for 168 hours (MEB versus PDB) and those of *A. yamanshiensis* (mannose versus fucoidan) for 21 days, at 25° C in the dark (static cultures in 6-well plates). Mycelia from each 6-well plate were collected in 50 mL polypropylene tubes (three replicates per strain and culture medium). Samples were then centrifuged for 15 minutes at 4° C and 4,000 rpm, filtered with Miracloth paper, freezed and homogenized in a mortar using liquid nitrogen, collected in 2 mL tubes and stored in liquid nitrogen until RNA extraction. RNA samples were obtained by using the NucleoSpin RNA Plant kit from Macherey-Nagel, following the protocol for RNA isolation from filamentous fungi and using RAP buffer for cell lysis. RNA samples were quality-checked by agarose electrophoresis (1.2%) and by measuring A260/A280 as well as A260/A230 ratios (NanoDrop 2000c, Thermofisher Scientific). Finally, samples were stored at -80° C.

Sequencing libraries were prepared with VAHTS Universal V10 RNA-seq Library preparation kit for Illumina (Vazyme). The kit converts the messenger RNA (mRNA) into dual-indexed libraries. mRNA was captured from total RNA using VAHTS mRNA Capture Beads 2.0 kit (Vazyme). Oligo(dT) magnetic beads purify and capture the mRNA molecules containing polyA tails. Input RNA was 100 ng of total RNA. The purified mRNA was fragmented and copied into first strand complimentary DNA (cDNA) by using reverse transcriptase and random primers. In a second strand cDNA synthesis step, dUTP replaced dTTP to achieve strand specificity. End-repair and d(A)-Tailing were combined with second strand cDNA synthesis step. The final step was ligation of adapters, using 1/10 dilutions of adapters. The resulting products were purified and size-selected for an insert size of 200-300 bp. The size-selected adapter-ligated fragments were amplified for sequencing on an Illumina system. The combination of both kits offered a polyA capture to selectively sequence mRNA, strand information on RNA transcripts, and unique dual (UD) indexing with unique combination of premixed Index 1 (i7) and Index 2 (i5) adapters (VAHTS Maxi Unique Dual Index DNA Adapters Set 1).

All libraries had an average size of 280-400 bp, with a mean size of 350 bp. Most of the samples showed a 140 bp peak which corresponded to adapter dimers. In order to remove the adapter dimers, all samples were pooled. An equimolar lane mix was prepared, calculated from the LabChip GX Touch profiles in the range of 200-100 bp, and a second round of bead purification was performed on the lane mix. Then, lane mix was quantified using QuBit dsDNA HS DNA kit. Library sequencing was carried out in an Illumina NextSeq 550 system (MidOutput Flow-cell; 150 cycles; paired-end; 2 x 76 bp read length). FASTQ files were demultiplexed and split into samples specific files (one per direction) using “Generate Fastq” module from NextSeq 550 software. All data processing and analysis was done in the Illumina BaseSpace Sequence Hub, using Dragen toolkits. Quality control of raw fastq files was carried out with Dragen FastQC + MultiQC tool v3.9.5 toolkit. Adapter and quality trimming and sequence length filtering was carried out with Dragen fastq v1.3.1 toolkit (minimum read length of 50 bp). Dragen reference builder v4.3.6 toolkit was used to build custom reference genomes for *A. yamanshiensis* and *M. marquandii*, while Dragen RNA v4.3.7 toolkit was used for alignment and mapping against custom reference genomes, and for differential expression analyses. Principal component analysis (PCA) plots were also generated with ClustVis [48] and plots for GO enrichment analyses and correlation matrices with R Studio (v2024.09.1+394; clusterProfiler package). See data availability in section 9.

### 2.8. Crude protein extracts and mass-spectrometry analyses

The same culture procedure as described in the previous section was followed for crude protein extraction from *A. yamanshiensis*. Samples (three replicates per carbon source: mannose or fucoidan from *F. vesiculosus* or from *U. pinnatifida*) were lyophilized overnight before protein extraction. Lyophilized samples were homogenized (Bullet Blender; Next Advance) and cultured in 1 mL of Drubin buffer [49] for 90 minutes at 4° C in a rotary shaker, before centrifugation at 14000 rpm and 4° C. Supernatants (crude protein extracts) were collected into new tubes. Aliquots of 100 µL per sample were preserved for protein quantification while the remaining extracts were stored at -80° C. Protein concentration was determined by measuring OD_595nm_ using Bradford reagent.

For in-solution tryptic digestion, proteins were first extracted by incubating for 30 minutes, at room temperature under agitation, the samples in a buffer containing 7 M urea, 2 M thiourea, and 4% CHAPS (3-[(3-Cholamidopropyl)dimethylammonio]-1-propanesulfonate hydrate). Digestion was carried out by following, with minor modifications, the FASP protocol described in [50]. Trypsin was added in 50 mM ammonium bicarbonate to a trypsin-to-protein ratio of 1:10, and the mixture was incubated overnight at 37° C. Peptides were dried out in an RVC 2-25 speedvac concentrator (Christ) and resuspended in 0.1% formic acid (FA). Peptides were desalted and resuspended in 0.1% FA using C18 stage tips (Millipore) prior to acquisition.

Proteomics analysis of the samples was carried out in a nano-liquid chromatography with electrospray ionization (nLC-ESI) equipment (Evosep One) coupled to trapped ion mobility/time of flight (TIMS/TOF) tandem mass spectrometer (timsTOF PRO, Bruker). Peptides from digested proteins were separated in a C18 nano-flow UPLC column (15 cm x 150 μm internal diameter, 1.9 μm particle size; EV1106, Evosep) using the Evosep 30SPD standardized method (44 minutes gradient). tims/TOF spectra of the samples were acquired through data independent acquisition (DIA). DIA data was processed with DIA-NN software for protein identification and quantification using default parameters [51]. Searches were done against *Albophoma yamanashiensis* M98 database (downloaded from https://mycocosm.jgi.doe.gov/Albya1) in library-free mode. Carbamidomethylation of Cys residues was considered as fixed modification and Met oxidation as variable modification. Data was loaded onto Perseus platform [52] for data processing (log2 transformation, imputation) and statistical analysis (Student’s t-test).

## 3. Results

### 3.1. Sordariomycetes, Dothideomycetes and Eurotiomycetes dominate cultivable fungi in coastal marine sediments of the Basque Country

A total of 273 isolates were obtained from coastal sediment samples (six estuarine stations and one coastal marine station; Figure 2A), exhibiting multiple phenotypic features (examples shown in Figure 2B). The phenotypes of 235 isolates (80% of the collection) were characterized in four of the culture media used for isolation (PDA, MEA, Pc and Lc; Figure S2) to identify isolates showing the same or similar phenotypes. Our results suggest that multiple isolates corresponded to the same strains, or strains of the same species showing subtle phenotypic differences (colored squares in Figure S2).

**Figure 2:**
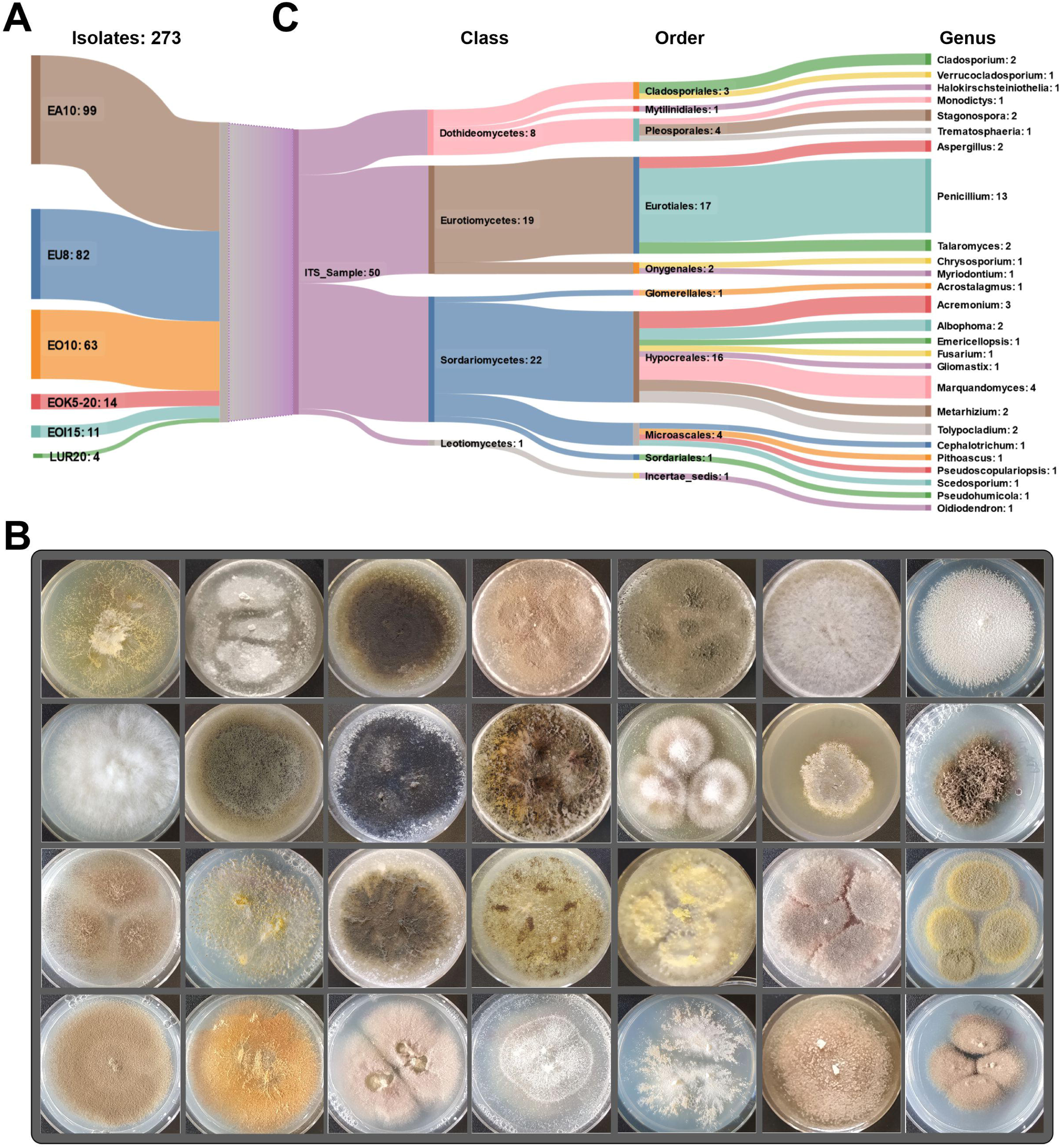
Phenotypic characterization of the library of filamentous microbes isolated from sediments collected along the Basque coast. The Sankey diagram shows A) the number of isolates obtained from each sediment sample (see Figure 1 and section 2), and B) the classes and genera represented in the subset of isolates whose ITS and/or *rpb1* regions were amplified (PCR) and Sanger-sequenced. C) Phenotypes of a random subset of isolates of our collection on the same culture medium (PDA, MEA, CMA, Pc or Lc) used for their isolation (see also Figure S2). The diameter of plates is 5.5 cm.

Genomic DNA of a number of isolates (50; 18% of the collection) was extracted. The ITS (internal transcribed spacer) region and/or an internal region of the gene *rpb1* were amplified by PCR from each DNA sample and sequenced (Figure 2C). Sordariomycetes (22) and Eurotiomycetes (19) were the most represented classes in the cultivable mycobiota of estuarine sediments of the Basque Country, followed by Dothideomycetes (8). *Penicillium* (Eutoriomycetes) was clearly the dominant genus (13), followed by *Marquandomyces* (4) and *Acremonium* (3) among Sordariomycetes and *Cladosporium* (2) and *Stagonospora* (2) among Dothideomycetes.

### 3.2. Potential pigment producers: Identification and phenotypic characterization of isolate M60 (*Marquandomyces marquandii*)

After a preliminary characterization of the isolates, we initially directed our search towards the identification of pigment producers (see examples in Figure 3A). Among pigment producers, isolate M60 (estuary of river Artibai, winter 2021) developed purple colonies and stained MEA and mainly PDA media in yellow (Figure 3B-C), indicating the secretion of pigments. Sanger sequencing of the ITS region suggested that isolate M60 was *Marquandomyces marquandii* [53], with a nucleotide identity above 99.4% when compared with the best NCBI hits GU566261.1 and OM647857.1.

**Figure 3:**
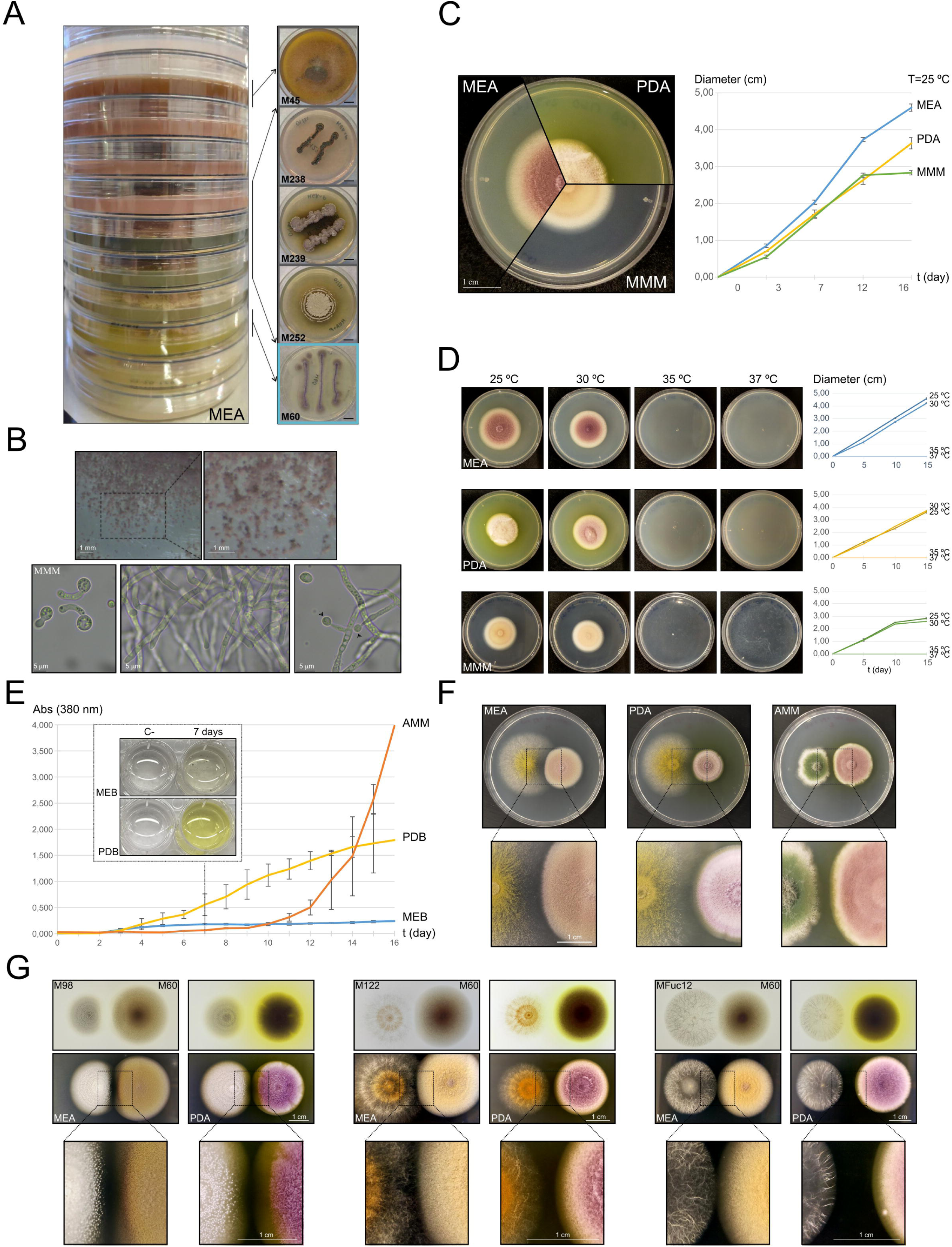
Phenotypic characterization of isolate M60 (*M. marquandii*). A) Lateral view of the culture plates (left) and phenotypes of some of the isolates (right) staining the culture medium due to the secretion of pigments, in MEA culture medium. The light blue square corresponds to isolate M60. B) Up: Developmental structures produced by *M. marquandii*. Scale bars: 1 mm. Down: Microscopy images showing germlings, mature hyphae and conidia. Scale bars: 5 µm. C) Phenotype and colony diameters of *M. marquandii* on MEA, PDA and MMM media after 0, 3, 7, 12 and 16 days of culture at 25° C. Values are calculated as the average ± standard deviation of three replicates per culture medium. Scale bar: 1 cm. D) Phenotype and colony diameters of *M. marquandii* on MEA, PDA and MMM media after 0, 5, 10 and 15 days of culture at 25°, 30°, 35° and 37° C. Diameter of Petri plates is 5.5 cm. Values on the graphs are calculated as the average ± standard deviation of three replicates per culture medium and temperature. E) OD_380nm_ values for the supernatants of *M. marquandii* cultures, measured daily (16 days) as the average ± sd of three replicates. Cultures were grown at 25° C. F) Phenotypes (up) and contact zones (down) between *M. marquandii* and *A. nidulans* on MEA, PDA and MMA media at 25° C. Scale bar: 1 cm. G) Phenotypes (top views in both rows 1 and 2) and contact zones (row 3) between *M. marquandii* and isolate M98 (*A. yamanashiensis*; left), isolate M122 (*genus Acrostalagmus*; middle) and isolate MFuc12 (*genus Gliomastix*; right) on MEA and PDA media at 25° C. See Experimental section for the time of inoculation of each species. Scale bar: 1 cm.

Microscopic analyses showed that M60 generated hyphal and developmental structures similar to those described for the species *M. marquandii* (Figure 3B) [53,54]. In liquid MMM and after 6 days of culture at 30° C and 180 rpm, we did not observe the formation of fully developed conidiophores but formation of side-branches in hyphae, each developing a single conidium [54]. The formation of these simplified asexual structures could be a consequence of carbon starvation, as occurs in the model fungus *Aspergillus nidulans* [55,56]. Radial growth of *M. marquandii* was bigger on MEA medium than on PDA or MMM medium (see the graph in Figure 3C). For example, after 16 days of culture at 25° C, the diameter of *M. marquandii* colonies was 4.60 ± 0.10 cm in MEA, 3.63 ± 0.15 cm in PDA and 2.83 ± 0.06 cm in MMM (n = 3 replicates in each culture medium). Growth was completely inhibited at temperatures equal or higher than 35° C (Figure 3D). We also followed coloring of growth media by measuring OD values at 380 nm for 14 days (three replicates per culture medium; Figure 3E). Pigmentation of the medium increased steadily in PDB medium (see the inset corresponding to day 7) and exponentially after day 10 in AMM.

Finally, the contact zones between *M. marquandii* and other fungal strains were analyzed in co-culture experiments. Spore pigmentation and colony diameter of *A. nidulans* changed drastically in MEA or PDA media (yellowish green colonies with bigger diameters) compared to AMM, where colonies were smaller, and conidia developed their characteristic dark green color (Figure 3F). Besides, the way in which colonies of both species contacted each other varied from MEA to PDA and mainly to AMM, with larger exclusion zones in the latter case. We also tested the colony phenotypes at contact zones between *M. marquandii* and specific strains from our collection of isolates (isolates M98: *A. yamanashiensis*; M122: genus *Acrostalagmus*; MFuc12: genus *Gliomastix*). In general, exclusion zones were larger when isolates were cultured on PDA compared to MEA (second row in Figure 3G), and this observation correlated with a more intense yellow pigmentation of the culture medium around *M. marquandii* colonies (first row).

### 3.3. Potential fungal degraders of fucoidan: Identification and phenotypic characterization of isolate M98 (*Albophoma yamanashiensis*)

Three types of fucoidan were used in the phenotypic screening, since the monosaccharide content, linkage type, degree of sulfation and the position of sulfate groups vary significantly among them [18,57–59]. Initially, we found that most isolates were able to grow with fucoidan from *L. japonica* (Figure S3), but this result may be biased due to impurities in this carbon source (purity of this polysaccharide not detailed on the supplieŕs website). In the case of fucoidan from *F. vesiculosus* or *U. pinnatifida* (purity above 95 %), radial growth of fungal isolates was significantly restricted compared to the use of fucoidan from *L. japonica* (Figures S3 and S4). In addition, *radii* of colonies of the isolates of our collection were, in general, higher when fucoidan from *U. pinnatifida* was used compared to that of *F. vesiculosus* (Figure S4). Isolate M98, isolated from the estuary of Artibai river in winter 2021, showed bigger radial growth in MMM supplemented with fucoidan of *F. vesiculosus* than in the negative control (Figure 4A), both when am monium tartrate and sodium nitrate were used as the nitrogen source (Figure S5). The same phenotype was observed for isolate M127, isolated from a sample of Oria river collected in summer 2021 (Figure S5). M98 showed a characteristic phenotype, producing pycnidia when cultured on PDA or MEA culture media (Figure 4B-C) [60,61]. These phenotypic traits had been previously described by Kobayashi and colleagues for *A. yamanashiensis* [62]. Thus, isolate M98 was initially associated to this hypocrealean fungus. Radial growth of isolate M98 was bigger on MEA and PDA media compared to MMM (Figure 4D). After 15 days of culture at 25° C, diameter of *A. yamanashiensis* colonies was 4.67 ± 0.06 cm in MEA, 4.50 ± 0.00 cm in PDA and 3.87 ± 0.12 cm in MMM (n = 3 replicates in each culture medium). Growth of this isolate was severely decreased at 35° C compared to 25° or 30° C, and impaired at 37° C.

**Figure 4:**
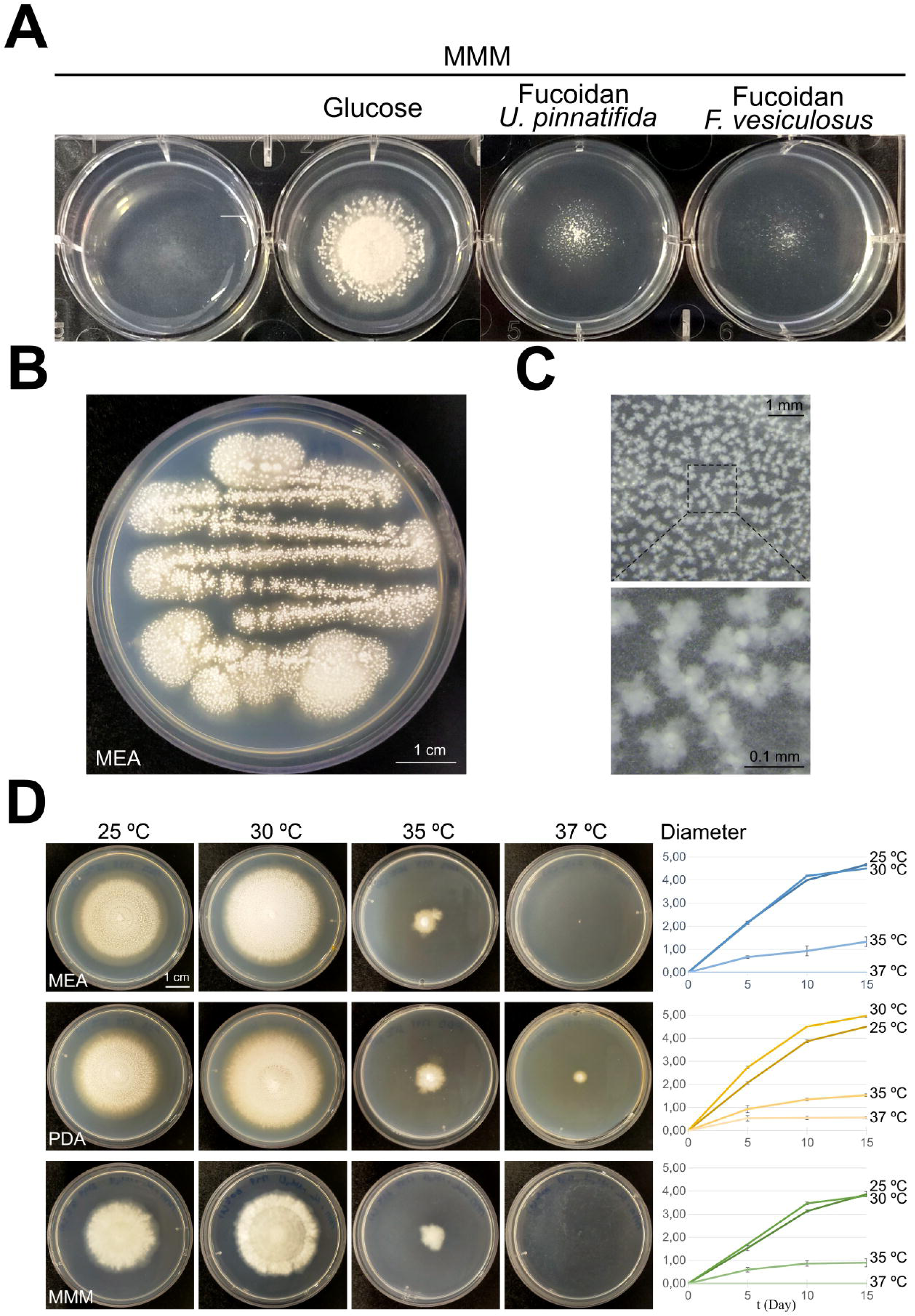
Phenotypic characterization of isolate M98. A) Phenotype on minimal medium supplemented with glucose or fucoidan from *Fucus vesiculosus* or *Undaria pinnatifida* (see Experimental section; see also Figure S5). B) Phenotype of M98 in MEA medium (scale bar: 1 cm), with the formation of pycnidia in panel C (scale bars, 1 and 0.1 mm, respectively). D) Phenotype and colony diameters on MEA, PDA and MMM media after 0, 5, 10 and 15 days of culture at 25°, 30°, 35° and 37° C. Diameter of Petri plates is 5.5 cm. Values on the graphs are calculated as the average ± standard deviation of three replicates per culture medium and temperature.

Sanger sequencing of the ITS region of isolate M98 indicated that this isolate was a species of the genus *Tolypocladium*, with a nucleotide identity above 99.8% in all cases (best hit NCBI accession number: AB457003.1). Alternatively, it may be identified as *Albophoma yamanashiensis*, with a nucleotide identity above 99.6% in all cases (best hit NCBI accession number: OW983802.1). Sequencing of *rpb1* suggested that it belonged to the genus *Tolypocladium* (nucleotide identity above 96.1%; best hit NCBI accession number: KF747240.1).

### 3.4. Genome sequencing and features of isolates M60 and M98

Isolate M60 was selected for genome sequencing due to its ability to stain the culture medium in yellow. Isolate M98 was selected due to its characteristic phenotype, including the apparent capability of growing on medium supplemented with fucoidan.

The size of the quality filtered assemblies of isolates M60 and M98 was 43.61 Mb and 31.18 Mb, respectively (see general data in Table 1; see their entry at Mycocosm: https://mycocosm.jgi.doe.gov/Marma1 and https://mycocosm.jgi.doe.gov/Albya1). The number of *contigs* longer than 2,000 bp were 97 and 38 for the two isolates. The largest *contigs* of M60 and M98 assemblies were of 5.62 and 4.47 Mbp, respectively. Annotation of each assembly led to the prediction of 12,408 and 10,216 proteins for isolates M60 and M98, respectively (Table 1). Completeness estimates were 97.2% for both assemblies and single-copy BUSCO shares ranged between 96.2% for M60 and 96.4% for M98, indicating high completeness and low redundancy of processed genomic data based on universal single-copy orthologs of Hypocreales. Average nucleotide identity (ANI) values were calculated among isolates and different species of the order Hypocreales (Figure 5A), including a reference genome of *A. yamanashiensis* (Genbank accession number BCKH01000001.1; Table S1). The highest ANI values for isolate M60 were obtained with species of *Metarhizium* and *Pochonia* (Figure 5A), which aligns with the results described by [53]. For isolate M98, the percent identity with the reference genome of *A. yamanashiensis* was 99%.

**Figure 5:**
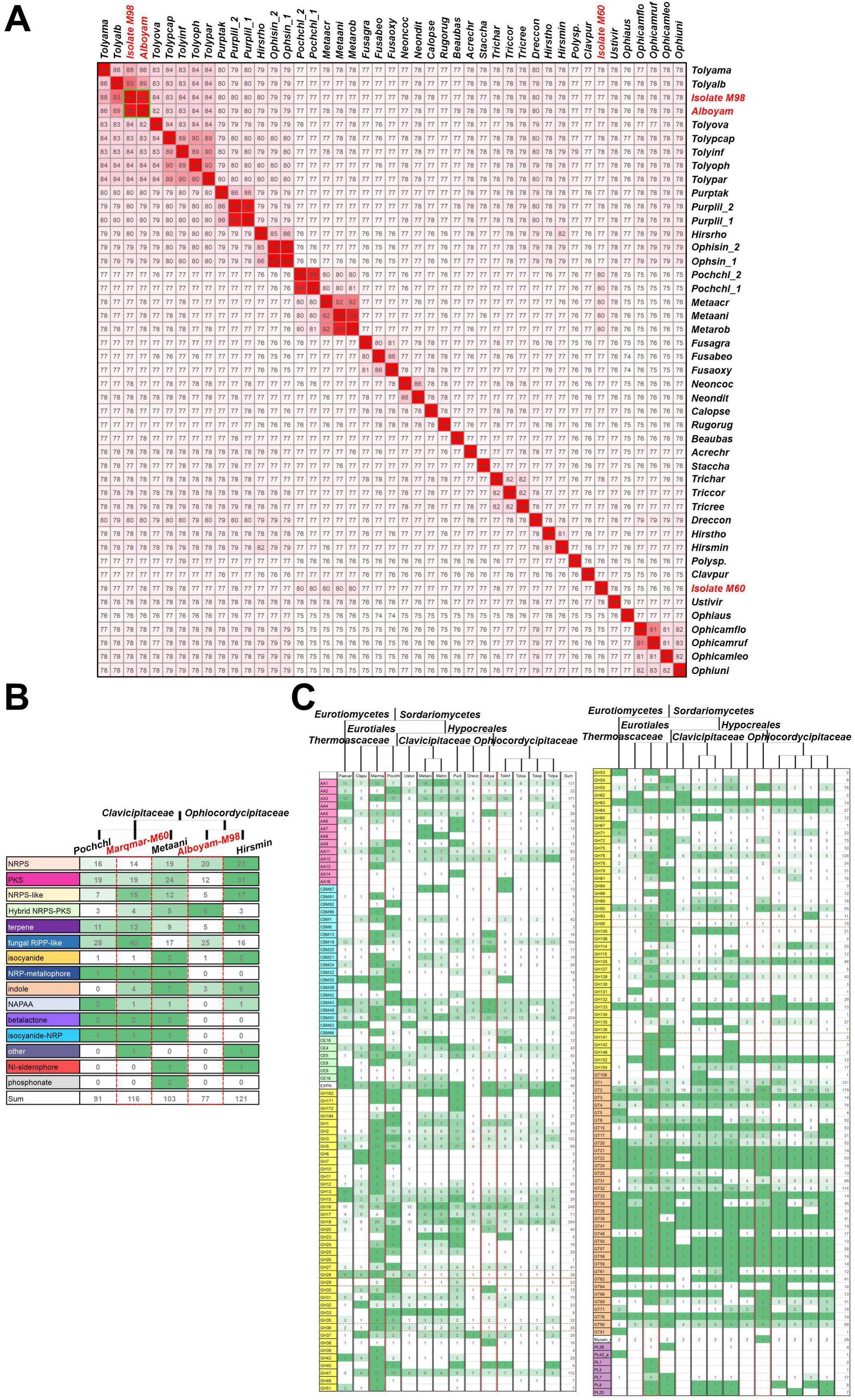
Genomics analyses of isolates M60 and M98. A) ANI analysis in which the genomes of M60 and M98 were compared to those of multiple Hypocreales species (see Table S1). B) Counts for genes encoding backbone enzymes of secondary metabolite gene clusters in the genomes of specific species of the order Hypocreales. The data in the heatmaps was retrieved from antiSMASH. C) Predicted CAZyme repertoires of isolates M60 and M98. The heatmap was generated using the data given by the MycoCosm annotation pipeline. Brown squares highlight GH families used by marine bacteria for fucoidan degradation [18]. For panels B and C, the trees in the heatmaps were generated with the common tree function at the NCBI taxonomy browser, using the IDs of the species analyzed. Red squares highlight data corresponding to *M. marquandii* and *A. yamanashiensis*.

**Table 1:**
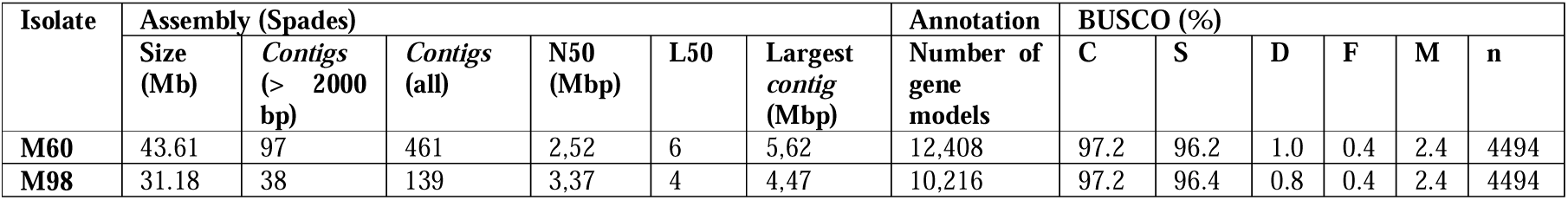
General data for genome sequencing analyses of isolates M60 and M98. C: Complete BUSCOs; S: Complete and single-copy BUSCOs; D: Complete and duplicated BUSCOs; F: Fragmented BUSCOs; M: Missing BUSCOs; n: Total number of BUSCOs for Hypocreales.

KOG (euKaryotic Orthologous Groups) analysis for isolates M60 and M98 were compared to those of Hypocreales species at MycoCosm. Since *Marquandomyces marquandii* was initially known as *Paecilomyces marquandii* [53,54], two species of the *genus Paecilomyces* (class: Eurotiomycetes; order: Eurotiales), *P. variotii* and *P. niveus*, were included in the comparison (Figure S6A). *M. marquandii* M60 showed the second highest content in KOG group Q (Secondary metabolites biosynthesis, transport and catabolism), while in the case of KOG groups G (Carbohydrate transport and metabolism) and M (Cell wall/membrane/envelope biogenesis) it was the first and third most abundant, respectively. In the case of KOG groups M and G, *A. yamanashiensis* was in both cases the sixth genome with the highest content. These values were in line with the total KOG counts for each species (second highest for *M. marquandii* and eight for *A. yamanashiensis*).

### 3.5. Genomic analysis of the potential for secondary metabolite production of *M. marquandii* M60 and *A. yamanashiensis* M98

Among the 21 genomes analysed in the KOG analysis, that of *M. marquandii* isolate M60 had a high number of backbone enzymes of secondary metabolite gene clusters (57, fourth genome in the ranking), while *A. yamanshiensis* M98 had 44 (seventh in the ranking; Figure S6B). However, the latter genome was particularly enriched in genes encoding hybrid NRPS (non-ribosomal peptide synthetase)-PKS (polyketide synthase) enzymes and NRPS enzymes, being at the second position of the ranking for both gene categories. In contrast, *M. marquandii* M60 had the second highest number of NRPS-like enzymes.

Considering the potential of *M. marquandii* for secondary metabolite and pigment production, and that of *A. yamanashiensis* as a producer of terpendole E, an inhibitor of enzymes involved in cholesterol metabolism [63], we annotated secondary metabolite gene clusters in the genomes of isolates M60 and M98 [45] by using antiSMASH (Figure 5B). We further compared this analysis with that at the MycoCosm website. The analysis confirmed previously observed general trends and suggested that the genomes of *A. yamanashiensis* and mainly *M. marquandii* encoded a high number of fungal RiPP-like compounds [64]. RiPPs are synthesized from precursor peptides in which a leader peptide is identified and removed by modifying enzymes, leaving a core peptide on which post-translational modifications occur [65].

AntiSMASH predicted the presence of 84 secondary metabolite gene clusters in the genome of isolate M60 (Figure S7), including 11 secondary metabolite gene clusters with similarity equal to or higher than 50%. Similarity of 100% was assigned to clusters c2.6 (ε-Poly-L-lysine, a non-ribosomal peptide), c3.3 (choline; non-ribosomal peptide), c9.1 (FR901512; polyketide), c15.1 (1,3,5,8-tetrahydroxynaphthalene; polyketide), and c16.1 (dymethylcoprogen, non-ribosomal peptide), while similarity percentages ≥80% was assigned to the clusters for the synthesis of sorbicillins and hypothemycin polyketides, both molecules with therapeutic activity as antifungals, antiparasitic and antitumor [66–68].

In the case of the *A. yamanashiensis* isolate (M98), antiSMASH predicted the presence of 59 secondary metabolite gene clusters, including 7 with similarity higher than 70% compared to those known to synthesize the compounds as indicated in Figure S8. A similarity of 100% was assigned to clusters c2.2 (terpendole E; terpene) [63], c5.2 (choline; non-ribosomal peptide), and c7.3 (polyketide + non-ribosomal peptide), the latter being necessary for the synthesis of tolypyridone C, a phenylpyridine with antibacterial and antifungal activity [69].

### 3.6. CAZyme repertoires of *M. marquandii* M60 and *A. yamanashiensis* M98

The predicted CAZyme (carbohydrate-active enzymes) repertoires of isolates *M. marquandii* M60 and *A. yamanashiensis* M98 were analyzed to assess their potential to degrade complex algal polysaccharides. A total of 656 and 403 proteins with CAZy domains were predicted for M60 and M98, respectively, by the CAZy annotation pipeline and available in MycoCosm (Figure 5C). Similarly, 610 (M60) and 365 (M98) CAZymes were predicted, respectively, by at least two of the tools used by dbCAN3 (Tables S2, S3). A total of 213 (34.9%; M60) and 143 (39.2%; M98) of the CAZymes predicted by dbCAN3 were also predicted to include a signal peptide in their sequence and would therefore represent the set of potentially secreted CAZymes of those isolates.

In the case of isolate M60, 101 proteins were predicted to include AA domains (auxiliary activity family protein) from CAZy classification, 47 proteins with CBM domains (carbohydrate-binding module), 23 with CE domains (carbohydrate esterase), 339 with GH domains (glycoside hydrolase), 123 with GT (glycosyl transferase) and 9 with PL (polysaccharide lyase) domains. For M98, dbCAN3 predicted a significantly lower number of GHs, 169, plus 53 AAs, 23 CBMs, 14 CEs and 121 GTs. Overall, the genome of *M. marquandii* M60 is predicted to encode the highest number of CAZymes among the species compared in this work. In most cases, this is probably associated with *M. marquandii* being the species with the highest number of predicted protein-coding genes. However, the genome of *Pochonia chlamydosporia* 170 is predicted to encode 14,204 proteins, compared to 12,408 in the case of *M. marquandii* M60 (1.14 times higher), while the number of predicted CAZymes in M60 is 649 compared to 538 in *P. chlamydosporia* (1.20 times higher in *M. marquandii* M60). *A. yamanshiensis* M98 is predicted to include a remarkable set of GT proteins, the highest number among the sequenced species of the Ophiocordycipitaceae family analyzed in Figure 5D. However, no PLs were identified, as occurs with sequenced genomes from the Clavicipitaceae family species *Claviceps purpurea* and *Ustilaginoidea virens* and the Ophiocordycipitaceae family species *Drechmeria coniospora* analyzed in this work.

Due to the ability of *A. yamanashiensis* isolate M98 to grow on medium supplemented with the recalcitrant polysaccharide fucoidan, we focused on the analysis of the CAZymes and sulfatases potentially involved in the degradation of this algal polysaccharide [18,70]. A limited number of proteins were found in M98, including one fucosidase of family GH29 and one of family GH95 (brown squares in Figure 5D). For comparison, 2 and 6 were predicted, respectively, in the case of *M. marquandii* isolate M60, plus 1 additional fucosidase of the GH141 family. Additionally, galactosidases and galacturonidases of GH36 (1) and GH28 (1) families were found in M98 (8 and 7, respectively, in the case of *M. marquandii* isolate M60).

Finally, SulfAtlas predicted that the genome of isolate M98 included genes encoding 7 sulfatases, of families S1_6 (2), S1_11 (1), S1_12 (1), S2 (1) and S3 (2). On the other hand, the genome of isolate M60 encoded 6 sulfatases, of families S1_4 (3), S1_8 (1) and S3 (2). Overall, the repertoire of predicted fucoidan-degrading enzymes and sulfatases in the fungal isolates characterized here is far more limited than the one described in the literature for fucoidan-degrading marine bacteria [18].

### 3.7. Comparison of the transcriptional patterns of *M. marquandii* M60 cultured on PDB versus MEB media: the sorbicillin gene cluster is strongly deregulated

To determine the set of activities induced in *M. marquandii* for the secretion of the yellow metabolite, we compared the transcriptomic response of mycelia cultured on PDB versus MEB culture media. Samples were collected, based on the graph in Figure 3E, after 7 days of static culture at 25° C in the dark. Both replicates of each sample grouped together, with high correlation values (Figure 6A-B). Reads were detected for 8,661 genes, 69.8% of the 12,408 genes annotated in the genome of *M. marquandii* M60. A total of 2,550 differentially expressed genes were identified (adjusted p < 0.05); 1,075 were significantly upregulated in PDB medium compared to MEB while 1,475 were downregulated (Figure 6C; Table S4). The heatmap in Figure 6D shows the correlation in the replicates for Top100 up- and downregulated genes. Gene enrichment and functional analysis of significantly deregulated genes and the proteins encoded by the Top20 up-/downregulated genes showed that processes such as transport, transcriptional regulation or oxidation-reduction processes were among the most significantly deregulated molecular functions (Figure 6E-F). Furthermore, we observed the presence of secondary metabolite gene cluster genes among the Top20 deregulated genes.

**Figure 6:**
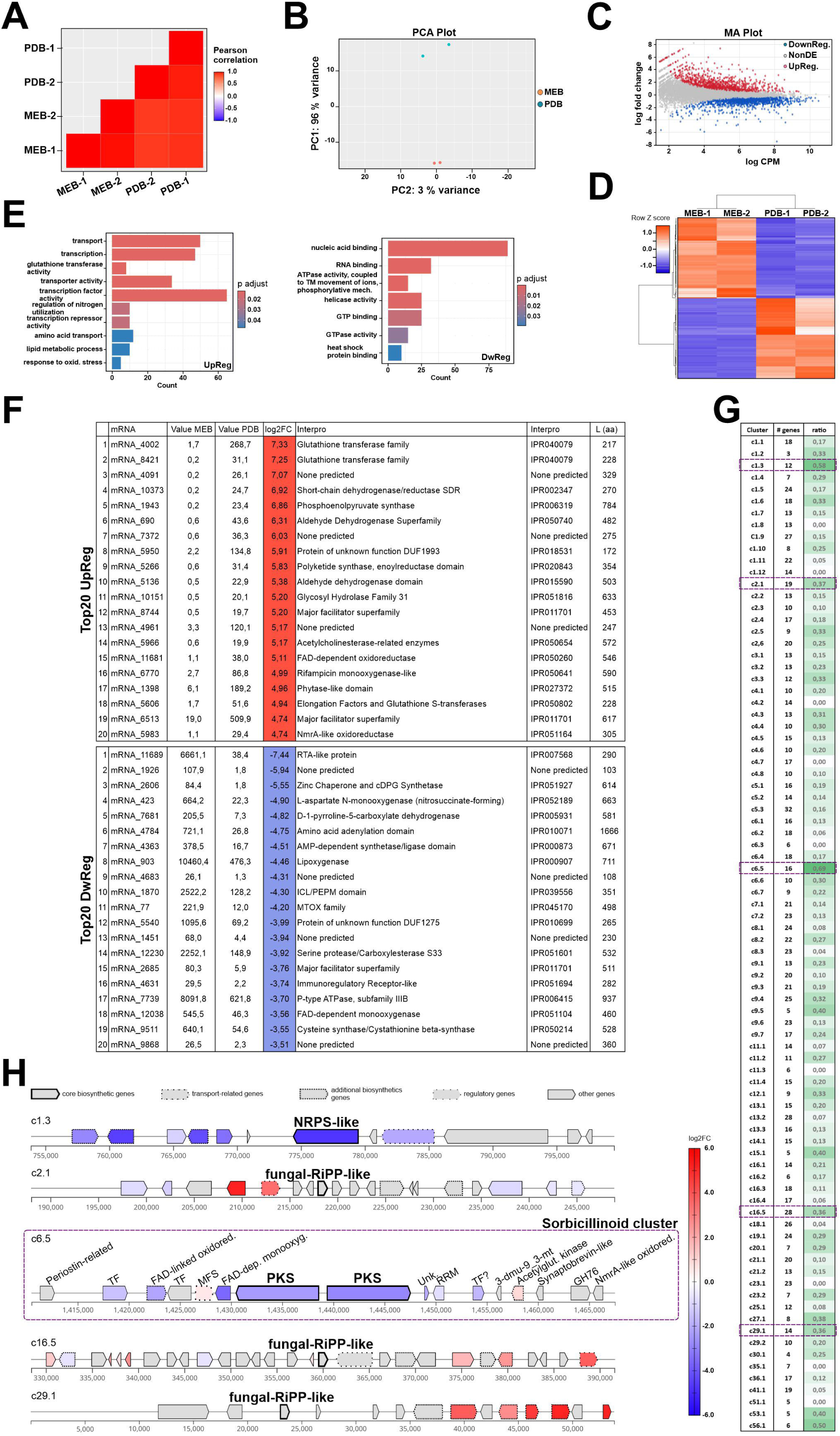
Transcriptional profiling of isolate M60 in PDB and MEB culture media. A) Correlation among the replicates (two per culture medium) analyzed. B) PCA plot for the replicates. C) MA plot showing genes up- and downregulated in PDB compared to MEB media. D) Heatmap showing the expression values for the Top100 up- and downregulated genes. E) GO enrichment analysis for genes up-(left) and downregulated (right) in PDB compared to MEB media. F) Functional analysis of Top20 up- and downregulated genes in PDB compared to MEB media. G) Ratios corresponding to the number of significantly deregulated genes in each secondary metabolite gene cluster and the number of cluster genes, in PDB versus MEB media. The list of genes from each secondary metabolite gene cluster were predicted by antiSMASH. Purple squares indicate clusters composed of more than ten genes and in which more than 35% of those genes were significantly deregulated in PDB compared to MEB. H) Gene organization in each of the secondary metabolite gene clusters highlighted in panel G. The color code corresponds to log2FC values. The predicted roles of the proteins encoded by cluster c6.5, predictably involved in the synthesis of a sorbicillinoid compound, are also included.

We compared the list of significantly deregulated genes (adjusted p < 0.05) with that predicted by antiSMASH for *M. marquandii* M60 (Figure 5C), which included 1,240 genes in 84 predicted clusters (an average of 15 ± 7 genes per cluster; 32 genes in the longest cluster, while the shortest one is predicted to be composed of 3 genes). We focused on long (> 10 genes) secondary metabolite gene clusters with at least 35% of those genes being significantly deregulated in PDB medium compared to MEB (purple squares in Figure 6G). Clusters c1.3 (0.58; 12 genes) and mainly c6.5 (0.69; 16 genes) were those with the highest ratio of significantly deregulated genes versus genes in that cluster (Figure 6G-H). According to antiSMASH, cluster c1.3 showed low similarity (20%) compared to the imizoquin gene cluster from *Aspergillus flavus* NRRL3357 [71]. Imizoquins are a family of tripeptide-derived alkaloids that are produced via a nonribosomal peptide synthetase-derived tripeptide and have a protective role against oxidative stress during *A. flavus* germination. Cluster c1.3 was downregulated, with the gene encoding the backbone enzyme being the Top6 among downregulated genes.

Cluster c6.5 was annotated by antiSMASH as the sorbicillinoid gene cluster of *M. marquandii* (87% of similarity compared to the sorbicillin cluster of *Acremonium chrysogenum* ATCC 11550; [68]). The characteristic yellow pigment of *M. marquandii* has been described as a nitrogen-containing sorbicillinoid (see Discussion) [72]. Although two genes within cluster c6.5 of *M. marquandii* M60, encoding an MFS transporter and an acetylglutamate kinase, respectively, are slightly upregulated, most genes are downregulated, with the two PKS-encoding genes of the cluster being, respectively, the Top73 and Top53, and a third gene the Top18 of downregulated genes (Figure 6H).

### 3.8. Transcriptional and protein profiles of isolate M98 in response to fucoidan

To identify the set of *A. yamanashiensis* M98 activities potentially involved in the processing of fucoidan from *F. vesiculosus* or *U. pinnatifida*, we compared its transcriptomics and proteomics responses to those in minimal medium supplemented with mannose, as a non-sulfated control monosaccharide that is present in some types of fucoidan. Mannose residues in these two types of fucoidan are less than 2% of the relative abundance of monnosaccharides [18]. Samples of *A. yamanashiensis* M98 were collected after 21 days of static culture at 25° C in the dark. The three replicates per condition grouped together, as can be seen in the PCA and correlation plots (Figure 7A-B). The following three comparisons were carried out: 1) genes differentially expressed in medium with fucoidan from *F. vesiculosus* versus mannose (FV_Vs_Mann), 2) genes differentially expressed in medium with fucoidan from *U. pinnatifida* versus mannose (UP_Vs_Mann), and 3) genes differentially expressed in medium with fucoidan from *U. pinnatifida* versus medium with fucoidan from *F. vesiculosus* (UP_Vs_FV). RNA-seq reads for more than 75% (75.3 to 77.1%) of the 10,216 genes annotated in the genome of *A. yamanashiensis* M98 were detected in all comparisons. A total of 2,476 genes were differentially expressed (adjusted p < 0.05) in the first comparison (1,137 upregulated and 1,339 downregulated in FV_Vs_Mann); 2,480 in the second comparison (1,099 up- and 1,381 downregulated in UP_Vs_Mann); and 2,801 in the third one (1,353 up- and 1,448 downregulated in UP_Vs_FV; Figure 7C; Tables S5-S7). Figure 7D (colored squares) highlight the presence of genes up- or downregulated only in medium with fucoidan from *F. vesiculosus*, only in fucoidan from *U. pinnatifida* or in both cases, suggesting that the transcriptional responses to the two types of fucoidan overlap partially, but there are also specificities in each response pattern (Figure 7E). This pattern can also be seen in the Top20 up- and downregulated genes (Figure 8) and, overall, probably reflects the structural and compositional similarities and divergences of both types of fucoidan [18] (see Discussion). Common to both comparisons (FV_Vs_Mann and UP_Vs_Mann) were 609 upregulated (53.6 and 55.4%, respectively) and 842 downregulated (62.9 and 61.0%, respectively) genes, while 528 (46.4%) and 497 (37.1%) genes were specifically up- and downregulated in the FV_Vs_Mann response, and 490 (44.6%) and 539 (39.0%) genes were specifically up- and downregulated in the UP_Vs_Mann response.

**Figure 7:**
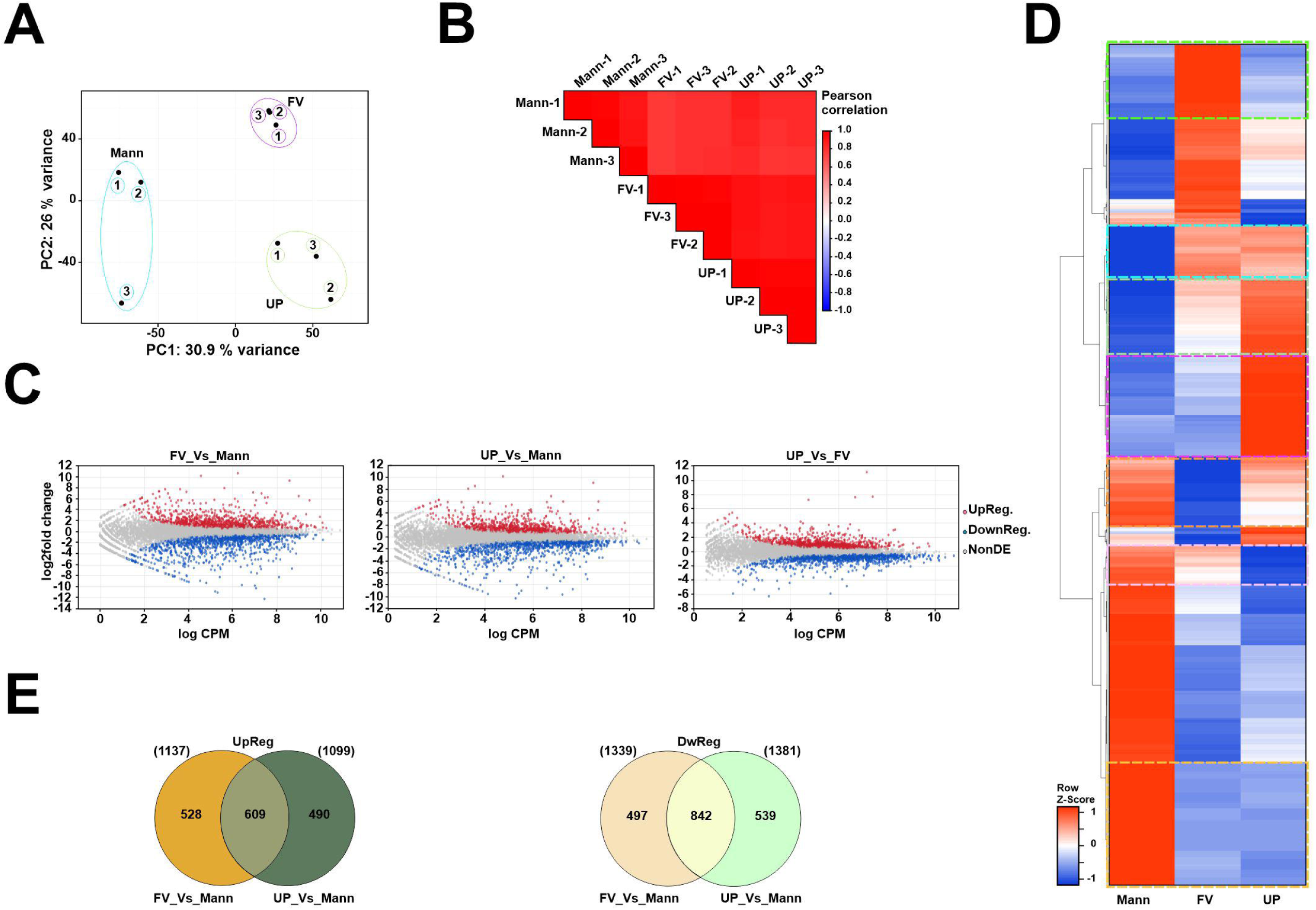
Transcriptional profilling of isolate M98 in culture medium supplemented with mannose or fucoidan. A) PCA plot for the nine samples sequenced. Mann: the three replicates for mycelia cultured with mannose as the main carbon source. FV: replicates for mycelia cultured with fucoidan from *F. vesiculosus*. UP: replicates for mycelia cultured with fucoidan from *U. pinnatifida*. B) Pearson correlation for the samples. C) MA plots showing genes up- and downregulated in three comparisons: FV Vs Mann, UP Vs Mann and UP Vs FV. D) Heatmap showing the expression patterns of genes differentially expressed in FV_Vs_Mann and UP_Vs_Mann comparisons. Multiple expression patterns are highlighted by the dotted squares. E) Venn diagram for up-(left) and downregulated genes in FV_Vs_Mann and UP_Vs_Mann comparisons, showing the number of genes differentially expressed in both comparisons or in one of them.

**Figure 8:**
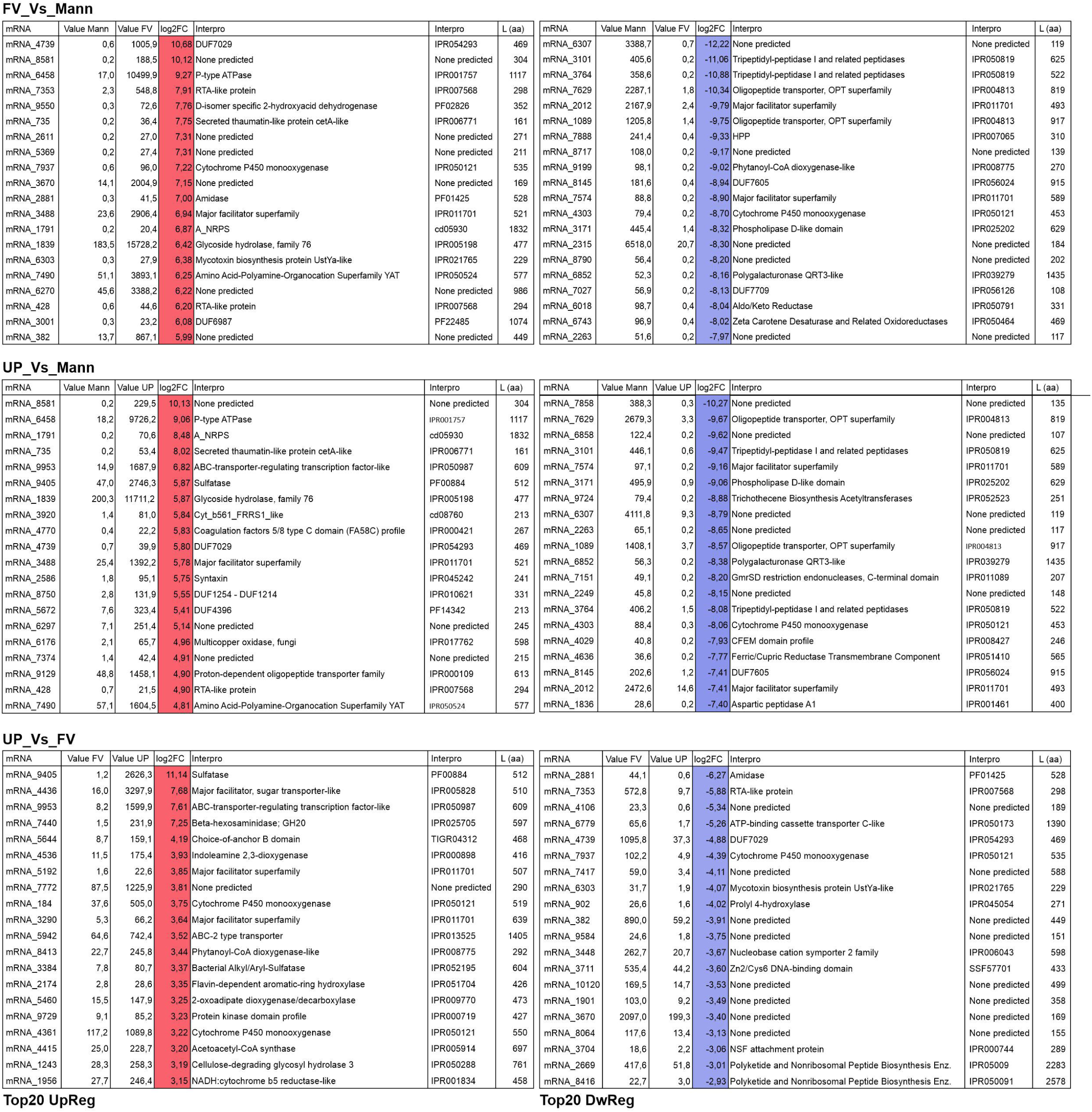
Top20 genes in each of the three transcriptomics comparisons carried out in this work with *A. yamanashiensis*. Count and fold change (log2FC) values are shown, together with the characteristic domains identified by InterPro.

The transcriptomics analysis was complemented with a proteomics analysis of crude protein extracts corresponding to samples collected under the same culture conditions. The three replicates per condition grouped together, as can be seen in the PCA and correlation plots (Figure 9A-B). The Heatmap in Figure 9C includes proteins with statistically significant changes (p < 0.05; Top300 of each comparison) in their levels in FV_Vs_Mann and UP_Vs_Mann comparisons (see also Table S8 with all proteomics data). It supports the trends in expression shown in the transcriptomics analysis in Figure 7D, strongly suggesting that *A. yamanashiensis* modulates, at least partially, the response depending on the type of fucoidan. Among the Top20 up- and downregulated proteins in FV_Vs_Mann and UP_Vs_Mann responses, the overlap was of 45.0% (up) and 15.0% (down), respectively (Figure 9D, top; see also Figure 10), while in the case of proteins with FV/Mann or UP/Mann ratios above 10 (up) and below 0.1 (down), the overlap was of 29%/43.9% and 29.2%/30.9%, respectively. We also compared the lists of differentially expressed genes and proteins (p < 0.05; Figure 9E for induced proteins; Figure S9 for proteins with lower levels). These lists matched partially, with a 23.3%/15.6% overlap in the case of the FV_Vs_Mann comparison, and 25.0%/18.7% overlap in the case of the UP_Vs_Mann comparison.

**Figure 9:**
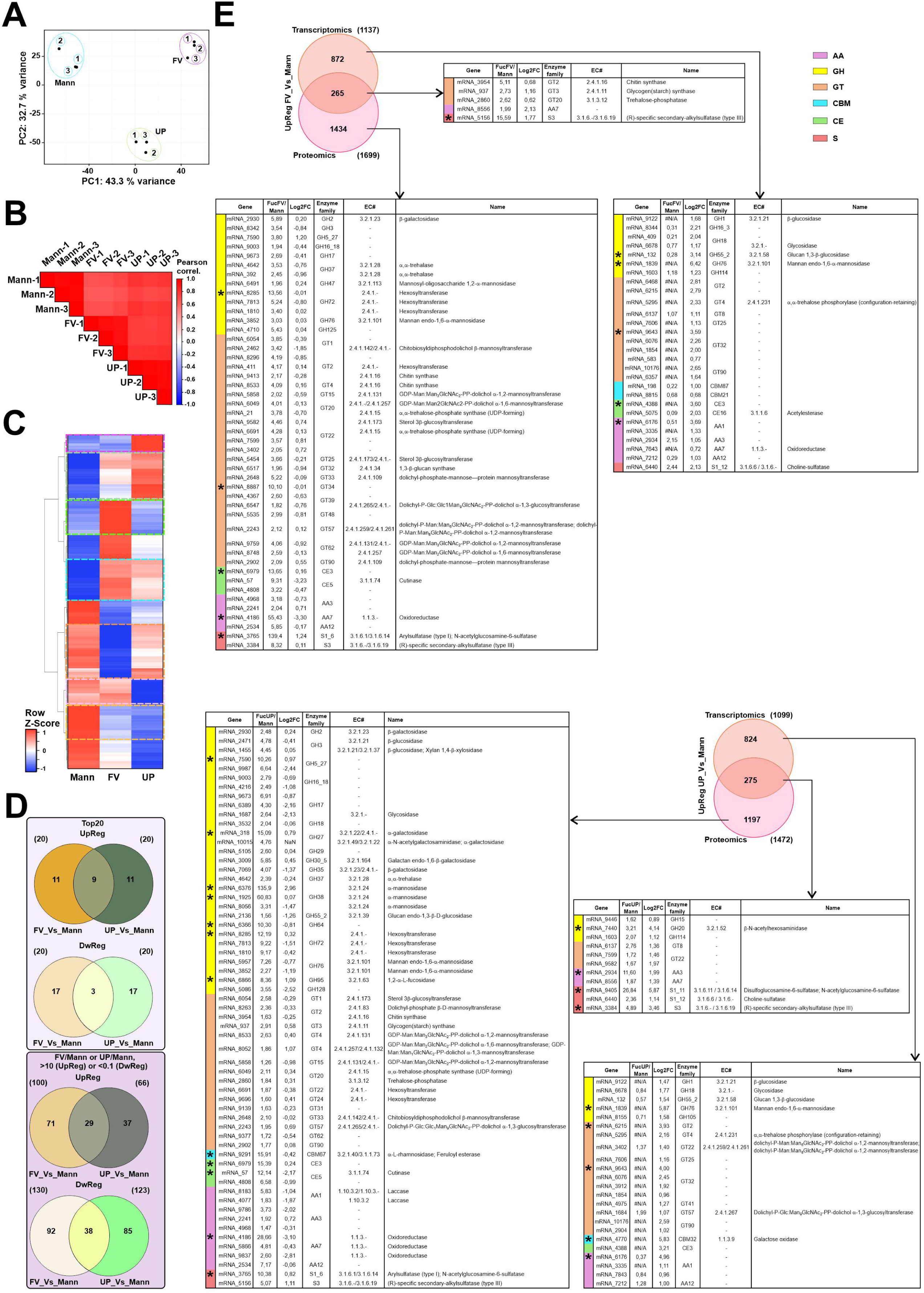
Comparison of transcriptomics and proteomics analyses of *A. yamanashiensis*. A) PCA plot for the nine crude protein samples sequenced. Mann: the three replicates for mycelia cultured with mannose as the main carbon source. FV: replicates for mycelia cultured with fucoidan from *F. vesiculosus*. UP: replicates for mycelia cultured with fucoidan from *U. pinnatifida*. B) Pearson correlation for the protein samples. C) Heatmap for proteins with statistically significant changes (p < 0.05; Top300 of each comparison, FV_Vs_Mann and UP_Vs_Mann) in their levels when isolate M98 is cultured on media supplemented with mannose or fucoidan from *F. vesiculosus* or *U. pinnatifida* as the carbon source. The dotted squares highlight groups of proteins with higher/lower levels only with fucoidan from *F. vesiculosus*, only with fucoidan from *U. pinnatifida*, or with both types of fucoidan. D) Venn diagrams showing the number of *A. yamanashiensis* proteins upregulated and downregulated in FV_Vs_Mann, UP_Vs_Mann or in both comparisons, when considering Top20 proteins or proteins with FV/Mann or UP/Mann ratios higher than 10 (UpReg) and below 0.1 (DwReg). E) CAZyme (dbCAN3) and sulfatase (SulfAtlas) analyses for the proteins and genes significantly upregulated in our proteomics and/or transcriptomics analyses. All genes and proteins (see Venn diagrams) with p values below 0.05 in the FV_Vs_Mann (top) and UP_Vs_Mann (bottom) comparisons were considered. Those proteins with ratios >10 and/or those genes with log2FC values >3 are marked with asterisks (an 1,2-α-L-fucosidase of the GH95 family is also highlighted). The same color code as in Figure 5D was used to designate CAZyme categories (plus S for sulfatases; in red). FV/Mann and UV/Mann ratios correspond to 2^exp^ ^(value^ ^FV−value^ ^Mann)^, 2^exp (value UP – value Mann)^ and 2^exp (value UP – value FV)^, respectively.

**Figure 10:**
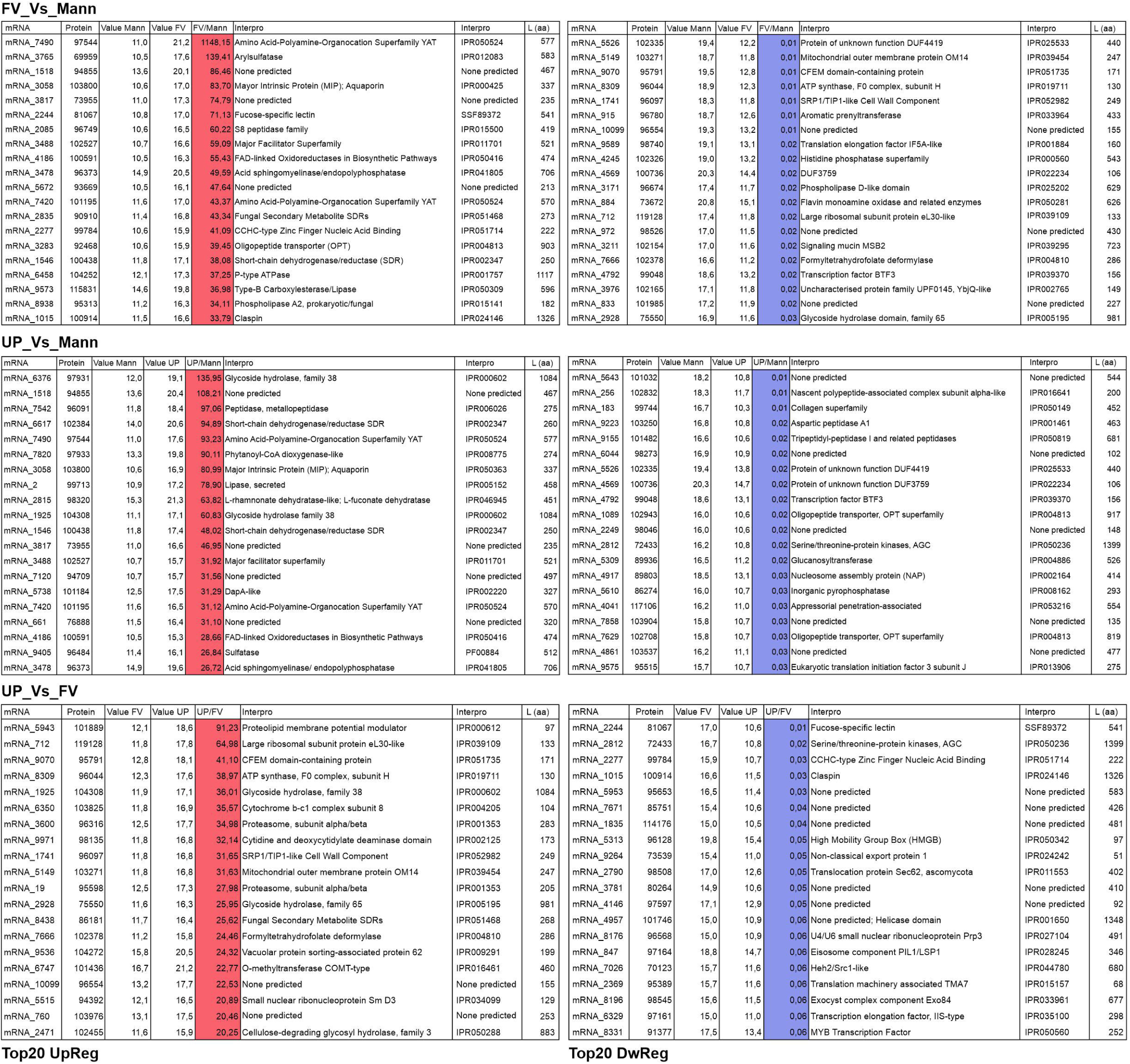
Top20 proteins in each of the three proteomics comparisons carried out in this work with *A. yamanshiensis*. FV/Mann, UV/Mann and UP/FV ratios correspond to 2^exp (value FV – value Mann)^, 2^exp (value UP – value Mann)^ and 2^exp (value UP – value FV)^, respectively.

We searched for CAZyme and sulfatases (and the corresponding genes) that were up- or downregulated at the protein level in the presence of fucoidan. In the FV_Vs_Mann comparison, one of the two S1_6 sulfatases (annotated as N-acetyl-D-glucosamine-6-sulfate 6-O-sulfohydrolase) of *A. yamanashiensis* was the Top2 protein with higher levels, with an FV/Mann ratio of 139.4 (Figure 9E and 10). Considering that sulfatases must be activated by sulfatase-modifying factors, we searched for genes encoding proteins with this domain (IPR005532) in the genome of *A. yamanshiensis*. One of them, mRNA_6814, was significantly upregulated in both FV_Vs_Mann (log2FC: 2.51) and UP_Vs_Mann (log2FC: 1.77) comparisons (statistically significant increase in protein levels only in the UP_Vs_Mann comparison). Furthermore, a protein annotated as a fucose-specific lectin was the Top6 (FV/Mann ratio of 71.1), a thiosulfate uptake-transporter was the Top38 (FV/Mann: 20.0), a sulfatase of family S3 ((R)-specific secondary-alkylsulfate sulfohydrolase) was the Top59 (FV/Mann: 15.6), and a glycosyl hydrolase of the GH72 family (β-1,3-glucanosyltransglucosidase) was the Top73 (FV/Mann: 13.6) protein (FV/Mann: 10.1) (indicated with asterisks in Figure 9E; see also Table S8). In the transcriptomics analysis, CAZyme-encoding genes with log2FC values higher than 3 (asterisks in Figure 9E) were predicted to belong to families GH76 (log2FC: 6.4; mannan endo-1,6-α-mannosidase), AA1 (3.7); CE3 (3.6; acetylesterase), GT32 (3.6; GDP-Man: α-1,3-mannosyltransferase), and GH55_2 (3.1; glucan 1,3-β-glucosidase).

In the UP_Vs_Mann comparison, two GH38s (mannosyl-oligosaccharide exo-α-1,2-mannosidase or mannosyl-oligosaccharide endo-α-1,P-mannosidase) were the Top1 and Top10 proteins, with UP/Mann ratios of 136.0 and 60.8, respectively. The Top19 protein (UP/Mann ratio of 26.8; log2FC of 5.9) was annotated by Sulfatlas as a S1_11 protein (exo-N-acetyl-D-glucosamine-6-sulfate 6-O-sulfohydrolase; Heparin/Heparan 6-O-sulfate sulfatase). In addition, Top21 and Top32 proteins were annotated as MFS-type L-fucose permeases (UP/Mann: 26.9 and 16.8, respectively), the Top34 as a protein containing a CBM67 domain (15.9; α-L-rhamnosidase), the Top37 as a protein of the GH27 family (15.1; α-galactosidase; galactomannan exo-α-1,6-galactosidase) and the Top47 as a GH72 (12.2; β-1,3-glucanosyltransglucosidase; GPI-anchored proteins that remodel β-1,3-glucans of the fungal cell-wall). The Top61 protein is the same S1_6 sulfatase that was the Top1 protein in the FV/Mann comparison (10.4; N-acetyl-D-glucosamine-6-sulfate 6-O-sulfohydrolase). Finally, the Top63 protein was annotated as a GH64 (10.3; glucan endo-β-1,3-glucosidase) and the Top64 as a GH5_27 protein (10.3; endoglycosylceramidase).

In the transcriptomics analysis, a gene annotated as encoding a sulfatase of the S3 family (log2FC: 3.5; (R)-specific secondary-alkylsulfatase, type III), a GH20 family protein-encoding gene (log2FC: 4.1; β-N-acetylhexosaminidase), the same GH76-encoding gene as in the FV_Vs_Mann comparison (log2FC: 5.9; mannan endo-1,6-α-mannosidase), a gene encoding a GT2-family protein (3.9; GDP-Man α-mannosyltransferase; α-1,4-L- or α-1,6-L-fucosyltransferase), the same GT32-encoding gene as in the FV_Vs_Mann comparison (4.0; GDP-Man: α-1,3-mannosyltransferase), a gene encoding a protein with a CBM32 domain (5.0; galactose oxidase), and a gene encoding a protein with an AA1 domains (5.0) were upregulated.

Overall, results from our proteomic and transcriptomic analyses for *A. yamanshiensis* strongly suggest that this fungus senses the presence of sulfated carbohydrates and activates a response mechanism (see Discussion).

## 4. Discussion

The application of the *dilution-to-extinction* method [11] in estuarine sediment samples enabled the isolation of a large diversity of filamentous fungi showing a variety of growth patterns. Our results suggest that cultivable fungi from the studied estuaries are mainly represented by Eurotiomycetes and Sordariomycetes species. This agrees with previous culture-based studies where marine fungi seemed to be dominated by Eurotiomycetes (*e.g.*, *Aspergillus* and *Penicillium*), Sordariomycetes (*e.g.*, *Fusarium*, *Trichoderma*, *Marquandomyces*) and Dothideomycetes (*e.g.*, *Cladosporium* and *Ulocladium*) (see references within [7]). Nevertheless, the extreme differences in growth rates among marine fungal species in culture [11] may be biasing the real composition of fungi in the samples, as typically found for bacteria (*i.e.*, the great plate-count anomaly; [73]). Additional efforts in isolation and culturing of slow-growing fungi and the direct identification of fungal taxa by metabarcoding (see, for example, [10,74,75]) will contribute to the identification of previously undescribed species, some of them potentially harboring biotechnological potential.

The repertoire of secondary metabolism backbone enzymes in fungi is composed mainly of PKS, NRPS, dimethylallyltransferases (DMATs), and terpene synthases [76], and the number of biosynthetic gene clusters per genome varies dramatically across species [76,77]. Here, we have focused on the description and comparison of specific genomic features of two species that were previously described phenotypically [53,54,62] but not at the genomic level. Interestingly, in both cases, the strains characterized phenotypically were originally isolated from terrestrial environments. On the one hand, *M. marquandii* has been reported to grow on mushrooms and soil, and their distribution covered Brazil, Netherlands, Russia, UK and USA [53]. On the other hand, *A. yamanshiensis* was isolated from a forest soil in Japan [62]. The isolation of strains of both species in sediment samples of Basque estuaries suggests that they are ubiquitously distributed and have adapted also to estuarine environments. While both isolates, M60 (*M. marquandii*) and M98 (*A. yamanashiensis*), correspond to species of the order of Hypocreales, the number of backbone enzyme-coding genes and secondary metabolite gene clusters is clearly higher in isolate M60 than in the genome of isolate M98, in correlation with the size of their genomes. However, the genome of isolate M98 is enriched in genes encoding specific backbone enzymes, such as hybrid NRPS-PKSs, compared to other Hypocreales species and includes gene clusters predictably involved in the synthesis of secondary metabolites of potential interest for biotechnology, such as Terpendole E, which acts as an acyl-CoA:cholesterol acyltransferase inhibitor [63] or as a kinesin Eg5 inhibitor [78].

There was a genome sequence of *M. marquandii* (synonims *Metarhizium marquandii* and *Paecilomyces marquandii*) available (Biosample: SAMN41475035), but the genome of this species had not been previously described in the literature. This fungus is able to produce a yellow pigment, which synthesis and secretion clearly depended on the composition of the culture medium (Figure 3A), as previously shown [53]. Using a *M. marquandii* strain isolated from a marine sediment sample collected in Argentina, Cabrera *et al.* described that the yellow pigment is a nitrogen-containing sorbicillinoid, sorbicillinoid urea, a Diels-Alder product of sorbicillinol [72,79]. In general, sorbicillinoids represent an important family of hexaketide compounds produced by terrestrial and marine fungi, including species of *Marquandomyces*/*Paecilomyces*, and have a variety of biological activities of interest in biotechnology including cytotoxic, antioxidant, anti-inflammatory, antiviral and antimicrobial activity [80,81]. Due to their biotechnological potential, recent studies have tried to develop fungal systems for sorbicillinoid hyperproduction [82]. Remarkably, we found that isolates closely related to M60, such as M43-45, affiliated with *Marquandomyces marquandii*, showed similar but not identical phenotypes. In the case of isolate M45, we have not observed the production of the characteristic yellow metabolite secreted by M60. Comparative genomic analysis of these strains, or mutagenesis screenings, could contribute to the identification of point substitutions or insertions/deletions causing an inhibition of the synthesis of the mentioned yellow pigment.

The gene cluster c6.5 in the genome of M60 is predictably involved in sorbicillin biosynthesis, as it shows high identity with the sorbicillin gene cluster of *Acremonium chrysogenum* (Figure 6H) [68]. In *Penicillium chrysogenum*, the sorbicillin biosynthesis gene cluster includes two PKS-encoding genes, *sorA* and *sorB*, and two transcription factor-coding genes, *sorR1* and *sorR2* (SorR1 acts as an activator, while SorR2 controls the expression of *sorR1*) [83]. Sorbicillin clusters have been described in additional fungal species, such as *Trichoderma reesei* [84]. Here, cluster c6.5 included genes encoding the two PKSs and the two transcription factors, and probably incorporates a gene encoding a third transcription factor (all three of the Zn_2_Cys_6_ family), adding a new layer of complexity to its regulation (Figure 6H). Our RNA-seq results suggest a general downregulation of the sorbicillinoid cluster of *M. marquandii* after 7 days of static culture at 25° C in PDB compared to malt extract broth, with the exception of the genes encoding an MFS transporter and a kinase. Figure 3E shows that at this time point the intensity of the yellow coloration of the PDB medium is still increasing. With these results, it is tempting to suggest that, in PDB medium, the concentration of the final product of the sorbicillinoid biosynthetic pathway of *M. marquanddi* is lower compared to the MEB medium and/or that the yellow pigment is not the final product of the biosynthetic pathway. However, the pigment would be accumulated in the PDB medium due to higher expression and protein levels of the transporter and a higher export capacity. Metabolomics analyses are needed to inform of the metabolic profile of organic extracts from *M. marquandii* grown on PDB or MEB media.

In the case of M98, this isolate was selected for genome analysis because the genome of *A. yamanshiensis* had not been described in detail and due to its apparent growth on fucoidan. This name refers to polysaccharides containing high percentages of L-fucose and sulfate ester groups, and are mainly derived from brown seaweed [70]. Fucoidans are highly diverse polysaccharides [18]. They can be grouped into homofucans (α-1,3-or alternating α-1,3-/α-1,4-linked fucose backbone with sulfate esters on positions *O*-2, *O*-3 or *O*-4) or hetero-polysaccharides with a non-fucose backbone consisting of mannose, galactose or glucuronic acid with side-branches of sulfated fucose [18]. These two types of main backbones can include additional modifications, increasing the diversity of fucoidans. The structure and composition of the two types of fucoidan used in this work were described previously by Sichert and colleagues [18]. That work also described that for fucoidan degradation bacteria use glycosyl hydrolases belonging to families GH29 (exo α-1,2-L-fucosidase), GH95 (α-L-fucosidase/α-L-galactosidase), GH141 (α-L-fucosidase/xylanase) and GH107 (endo-α-L-fucanase). The same authors also described that families GH36 (α-galactosidase, α-N-acetylgalactosaminidase), GH116 (β-glucosidase, β-xylosidase, acid β-glucosidase/β-glucosylceramidase, β-N-acetylglucosaminidase), and GH117 (β-D-galactofuranosidase) may be fucoidanases with a yet unknown function, and that families GH28 (polygalacturonase) and GH155 (glucuronidase; family removed from the CAZYme database; [85]) remove acetate, galacturonic acid and glucuronic acid, which are present in specific types of fucoidan. This set of GHs include both exo and endo activities, degrading internal and external (usually from the non-reducing end) glycosidic bonds. Regarding the analysis of sulfatases, subfamilies S1_15, S1_16, S1_17 and S1_25 have been reported as necessary for fucoidan degradation or enriched in fucoidan-degrading marine bacterial proteomes [18]. A recent work has shown that commercial fucoidan from *U. pinnatifida* (Sigma F8315) causes an upregulation of the same CAZyme and sulfatase families in Planctomycetota bacteria [86]. Considering that the same product reference was used in the present work and that the genome of *A. yamanshiensis* encodes a completely different repertoire of CAZyme and sulfatases, as seem to do fungi in general, the identification of the enzymes induced in response to fucoidans with different structures is interesting for biotechnology.

As a result, we have identified several induced sulfatases, such as those predictably belonging to families S1_6, S1_11 and S1_12. The genome of *M. marquandii* is not predicted to encode these sulfatase activities. For example, fucoidan from *U. pinnatifida* causes strong upregulation and higher protein levels of an S1_11 sulfatase. The Sulfatlas website associates this family with desulfation of polysaccharides such as heparin or chondroitin and glycoproteins such as mucins [87]. Thus, it will be interesting to determine in the future if fungal S1_11 sulfatases can act on sulfate groups of specific types of fucoidan. Furthermore, and considering that according to the Sulfatlas database only two experimentally determined structures of S1_11 sulfatases are available in the Protein Data Bank database (each with or without a ligand), both of them belonging to gut bacteria [88,89], this protein from *A. yamanshiensis* would constitute the first experimentally determined structure of a fungal S1_11 sulfatase.

In combination with sulfatases, the apparent induction of a sulfatase-modifying factor-coding gene, fucose-specific lectins, and L-fucose permeases strongly suggest that *A. yamanshiensis* senses and responds to the presence of carbohydrates composed of sulfated L-fucose. Although the set of upregulated GHs identified in our transcriptomics and proteomics analyses includes an exo-1,2-α-L-fucosidase (GH95), in general the CAZyme families point to processing of mannans and glucans. These two polysaccharides are components of the fungal cell wall, and their relative abundance varies depending on the clade. For example, mannans are more abundant and essential in the outer cell wall of *Candida*, while they are much less abundant in *Aspergillus* [90,91]. Although the composition of the cell wall of *A. yamanshiensis* has not been determined, the induction of those activities points to a model in which this fungus responds by remodeling its cell wall structure, maybe to obtain a more easily metabolizable carbon source. Future experiments will have to determine whether these enzymes have hitherto undescribed activities on fucoidan or are employed by fungi to remove mannose from fucoidan, although the percentage of this monosaccharide is lower than 2% in the types of fucoidan used in this work [18]. However, it can be concluded from the analysis of fungal genomes that if fungi are able to process recalcitrant marine polysaccharides such as fucoidan, they follow completely different mechanisms compared to bacteria.

## 5. Acknowledgements

We want to thank all groups and members of the EMOTION consortium, funded by the Basque Government (see Funding), for their support. The authors acknowledge technical and human support provided by the General Research Services (SGIker) of the UPV/EHU (ERDF and ESF funding), especially Dr. I. Bernales and Dr. I. Miguel, and the CNAG-CRG for assistance with genome and transcriptome sequencing projects.

## 8. Author contributions

O.E. and L.A.-S. conceived the project. A.O., Z.A., C.P.-C, R.L., M.D., M.O., R.L. and O.E. performed the experiments. M.A. and F.E. carried out proteomics analyses. A.L. assembled the genomes of *M. marquandii* and *A. yamanshiensis*, and S.H. and I.V.G. provided and coordinated annotation resources. All authors analyzed the results. O.E. and Z.A. wrote the first version of the manuscript. All authors edited and improved the manuscript.

## 9. Availability of Data and Material

The genome assemblies and annotations for *Marquandomyces marquandii* M60 and *Albophoma yamanshiensis* M98 are available from JGI MycoCosm (https://mycocosm.jgi.doe.gov/Marma1 and https://mycocosm.jgi.doe.gov/Albya1) and have also been deposited at DDBJ/ENA/GenBank under accessions **JAUIRS000000000** (BioProject **PRJNA990752**; BioSample **SAMN36274215**) and **JAUIRT000000000** (BioProject **PRJNA990765**; BioSample **SAMN36274349**), respectively. RNA-seq data for *M. marquandii* (MEB vs PDB media) were deposited under BioProject **PRJNA1233808** and BioSample **SAMN47279085** accession codes. RNA-seq data for *A. yamanashiensis* (mannose vs fucoidan) were deposited under BioProject **PRJNA1233821** and BioSample **SAMN47279930** accession codes.

## Supplementary Figures

**Figure S1:** Phenotype of a subset of isolates of our library in minimal medium with decreasing concentrations of bacteriological agar, without (left column) and with (right column) glucose (2%). Considering the phenotype of the strains and solidification of the agar, a 0.5% concentration was selected for the subsequent phenotypic characterization of the isolates.

**Figure S2:** Phenotypes of a subset of 235 isolates of our library in PDA, MEA, Pc and Lc culture media, all supplemented with penicillin, grown at 25° C in the dark. Squares highlight isolates probably corresponding to the same strain (*e.g.*, isolates M106, M107 and M108) or different strains of the same species (*e.g.*, isolates M43, M44, M45 and M60).

**Figure S3:** Phenotype of selected isolates on plates filled with minimal medium supplemented with glucose, laminarin, alginate or fucoidan from *Laminaria japonica*, or without supplementation of any of these carbohydrate sources (negative control).

**Figure S4:** Phenotype of selected isolates on plates filled with minimal medium supplemented with 2 g/L of glucose or fucoidan from *Fucus vesiculosus* or *Undaria pinnatifida*, or without supplementation of any of these carbohydrate sources (negative control).

**Figure S5:** A) Phenotype of isolates M98 and M127 on minimal medium supplemented with laminarin or fucoidan from *Fucus vesiculosus* or *Undaria pinnatifida*, compared to the phenotype on MEA medium, MMM or MMM supplemented with glucose. Ammonium tartrate and sodium nitrate were tested as the main nitrogen source. glc: glucose; lam: laminarin; fucoi: fucoidan. Six-well plates were used.

**Figure S6:** A) KOG analyses of the genomes of M60 and M98 in comparison with specific Hypocreales species. All the data in the heatmap was retrieved from MycoCosm. C) Counts for genes encoding backbone enzymes of secondary metabolite gene clusters in the genomes of specific species of the order Hypocreales (plus the Eurotiales *P. niveus* and *P. variotii*). The data in the heatmap was retrieved from MycoCosm. For both panels, the trees in the heatmaps were generated with the common tree function at the NCBI taxonomy browser, using the IDs of the species analyzed. Red squares highlight data corresponding to *M. marquandii* and *A. yamanashiensis*.

**Figure S7:** Table showing the results obtained in the analysis of the potential secondary metabolite gene clusters predicted in the genome of isolate M60 (*Marquandomyces marquandii*) by antiSMASH. The table includes the node the potential cluster is located in, the backbone enzyme(s) of each cluster, coordinates, the most similar known cluster, the type of metabolite predictably synthesized and the similarity with the reference cluster. Similarity, in this case, refers to the percentage of genes within the closest known cluster that have a significant BLAST hit to genes within the predicted cluster

**Figure S8:** Table showing the results obtained in the analysis of the potential secondary metabolite gene clusters predicted in the genome of isolate M98 (*Albophoma yamanashiensis*) by antiSMASH.

**Figure S9:** CAZyme and sulfatase analyses for the proteins and genes significantly downregulated in our proteomics and/or transcriptomics analyses. All genes and proteins with p values below 0.05 in the FV_Vs_Mann (top) and UP_Vs_Mann (bottom) comparisons were considered (see Venn diagrams). Those proteins with ratios <0.1 and/or those genes with log2FC values <-3 are marked with asterisks. The same color code as in Figure 5D was used to designate CAZyme categories (plus S for sulfatases).

**Supplementary Tables**

**Table S1:** Genomes used as reference in ANI analyses. GenBank and BioSample accession numbers are included.

**Table S2:** Predicted CAZYme repertoire for isolate M60. The analysis was carried out at the dbCAN website (Zhang *et al.*, 2018), and the table includes the CAZYme family each protein is linked to, the enzyme commission (EC) number, the tools that predicted this association as well as the hypothetic presence of a signal peptide in each predicted CAZYme.

**Table S3:** Predicted CAZYme repertoire for isolate M98.

**Table S4:** Differentially expressed genes of *M. marquandii* in PDB versus MEB media.

**Table S5:** Differentially expressed genes of *A. yamanashiensis* in medium supplemented with fucoidan from *F. vesiculosus* versus mannose as the main carbon source.

**Table S6:** Differentially expressed genes of *A. yamanashiensis* in medium supplemented with fucoidan from *U. pinnatifida* versus mannose as the main carbon source.

**Table S7:** Differentially expressed genes of *A. yamanashiensis* in medium supplemented with fucoidan from *U. pinnatifida* versus fucoidan from *F. vesiculosus*.

**Table S8:** Proteins of *A. yamanashiensis* with statistically significant changes in their levels when the minimal culture medium was supplemented with: 1) fucoidan from *F. vesiculosus* as the main carbon source compared to a medium supplemented with mannose; 2) fucoidan from *U. pinnatifida* as the main carbon source was compared to a medium supplemented with mannose; and 3) fucoidan from *U. pinnatifida* as the main carbon source was compared to a medium supplemented with fucoidan from *F. vesiculosus*. FV/Mann ratios correspond to 2^exp^ ^(value^ ^FV−value^ ^Mann)^, UP/Mann ratios correspond to 2^exp^ ^(value^ ^UP−value^ ^Mann)^, and UP/FV ratios correspond to 2^exp^ ^(value^ ^UP−value FV)^.

