## Supplementary figures and images for "Phenotypic and *omics* analyses of the Sordariomycetes *Marquandomyces marquandii* and *Albophoma yamanashiensis* isolated from estuarine sediments"

### Figure S1

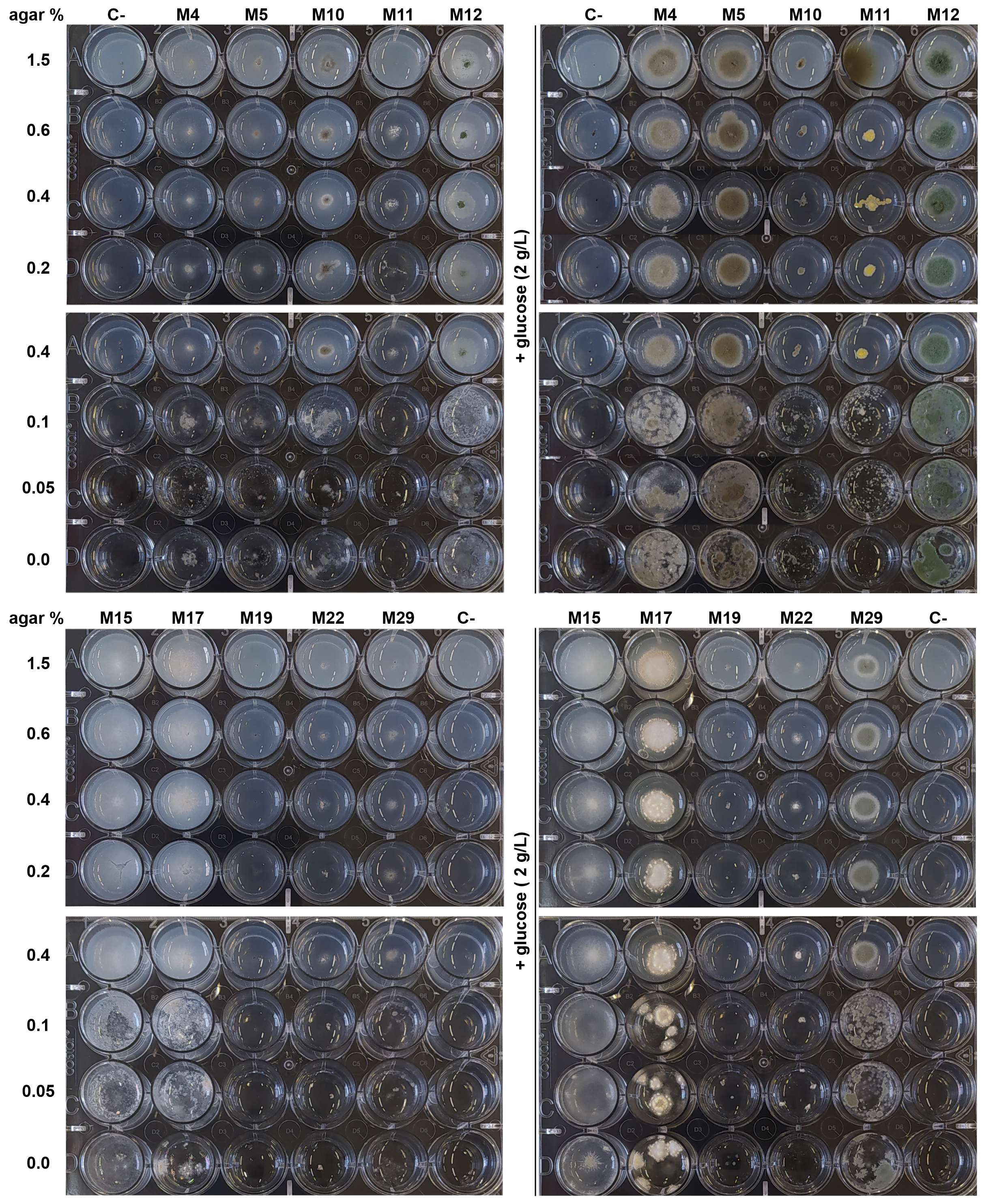

### Figure S2A

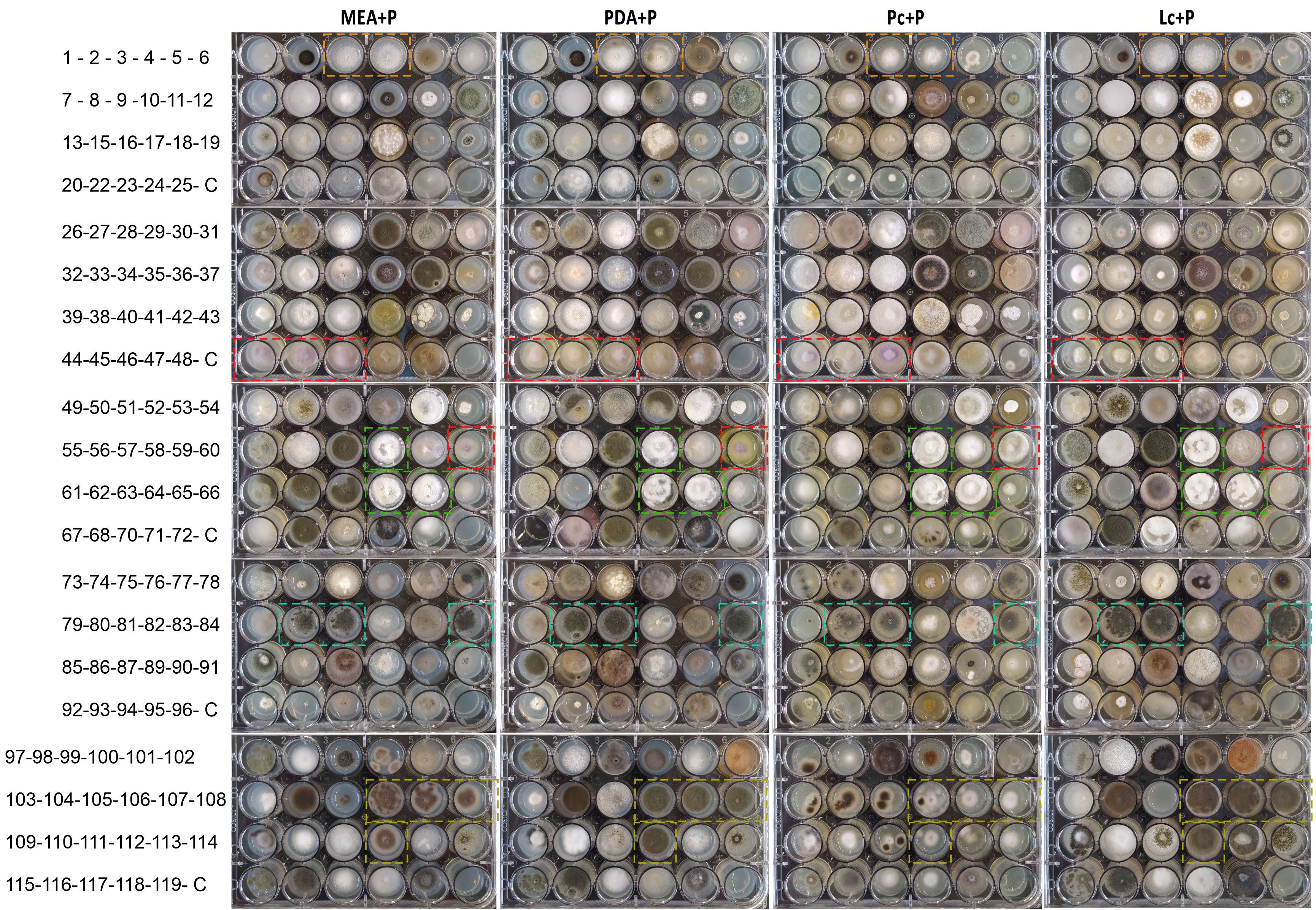

### Figure S2B

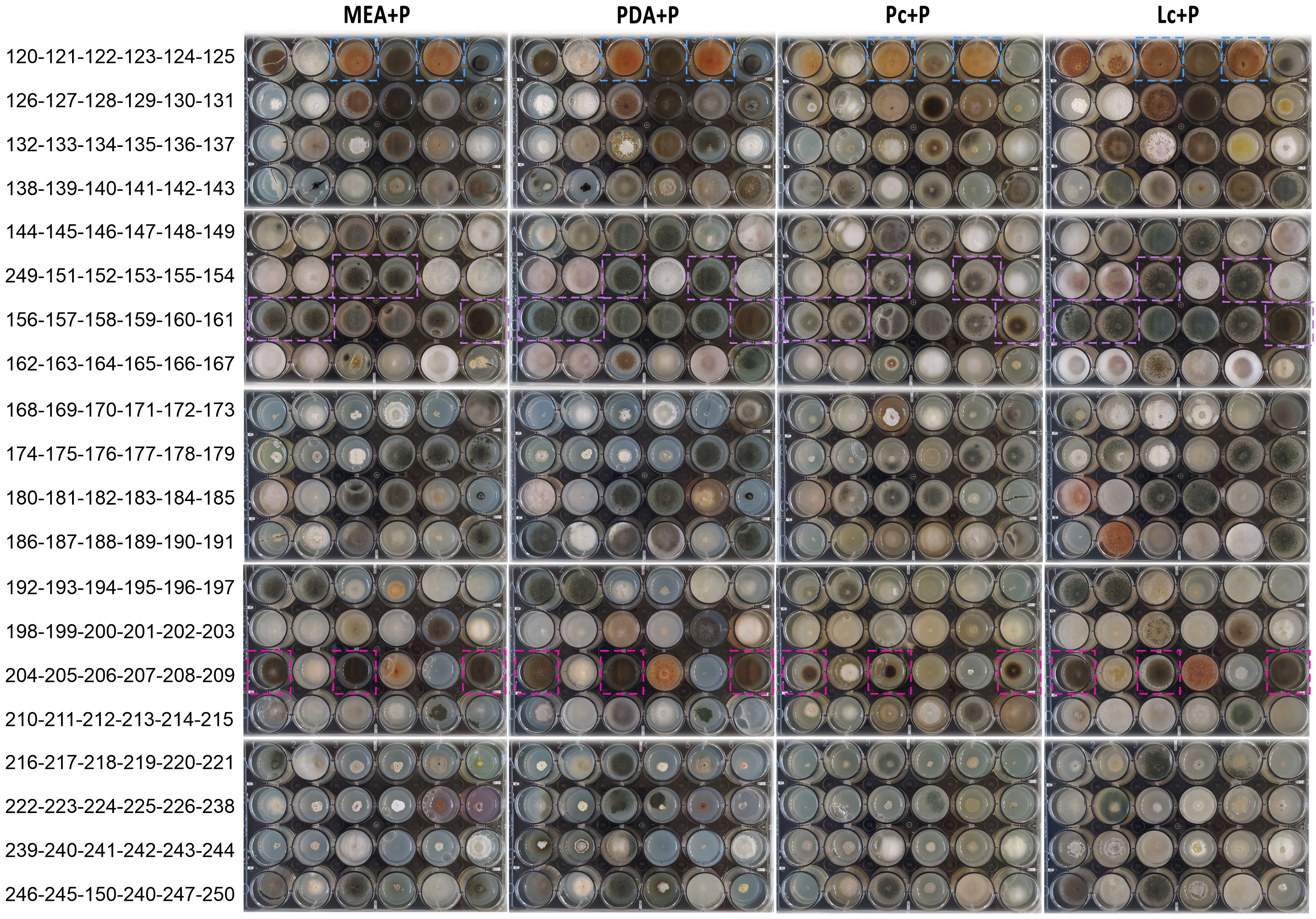

### Figure S3

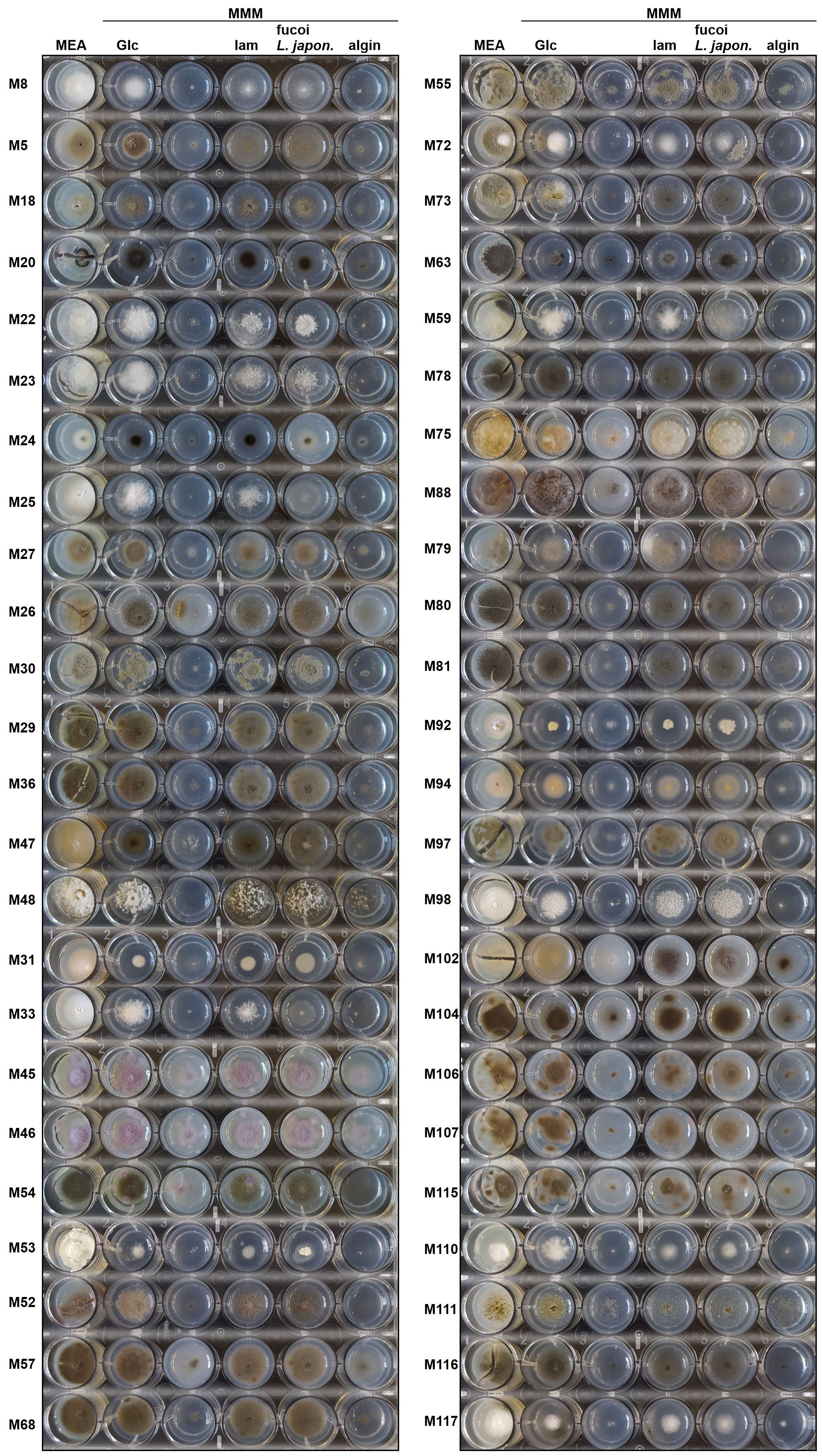

### Figure S4

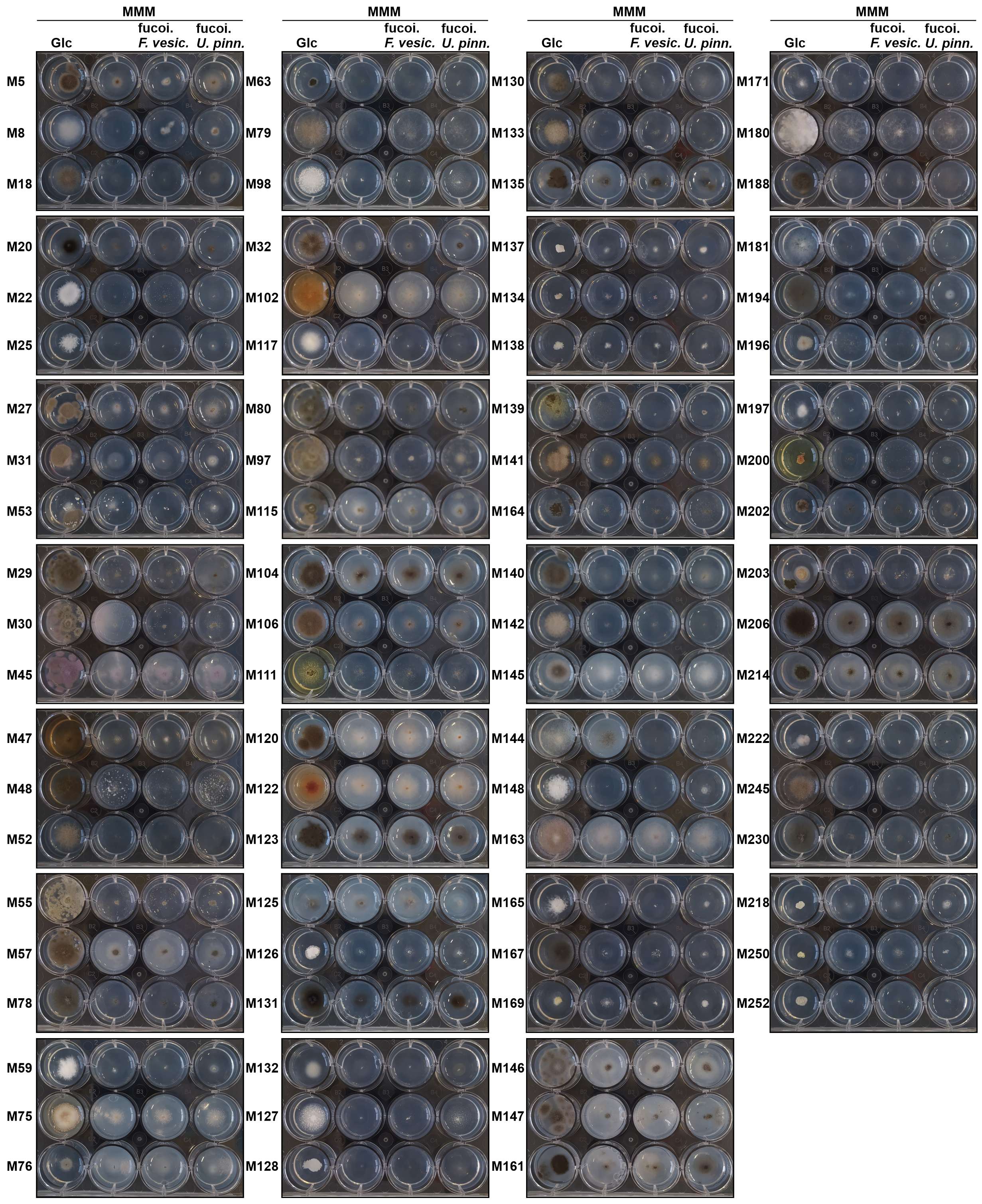

### Figure S5

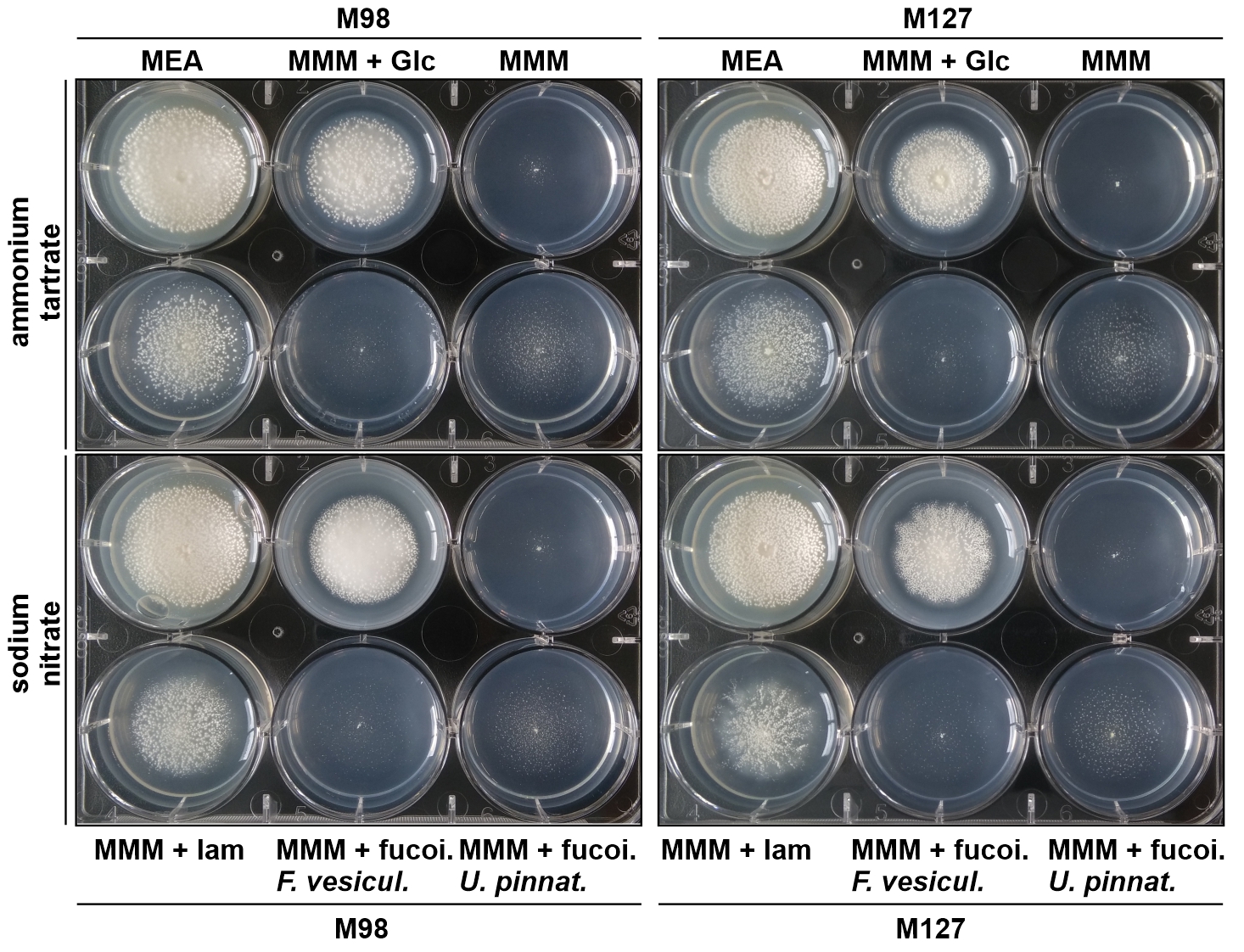

### Figure S6

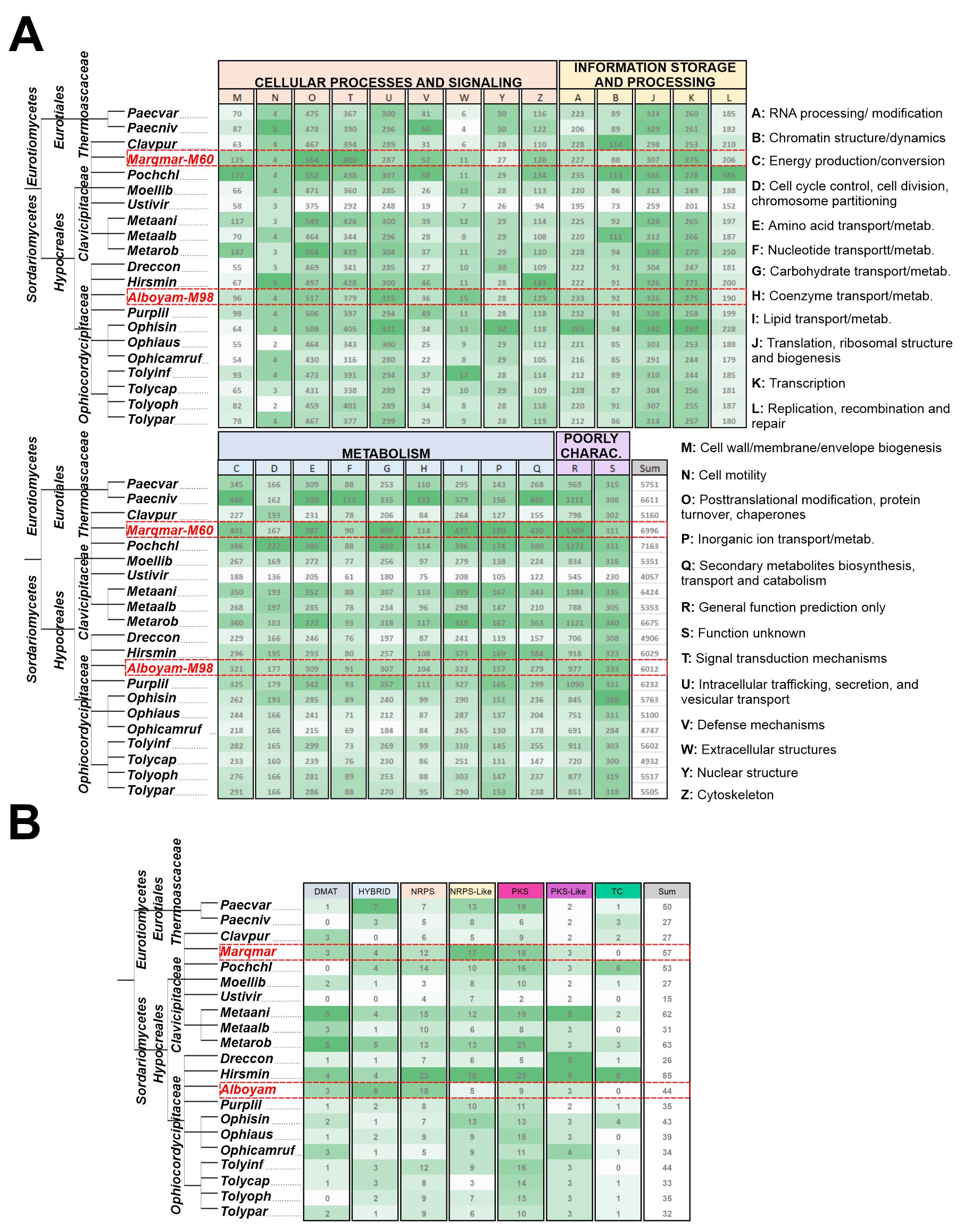

### Figure S7A

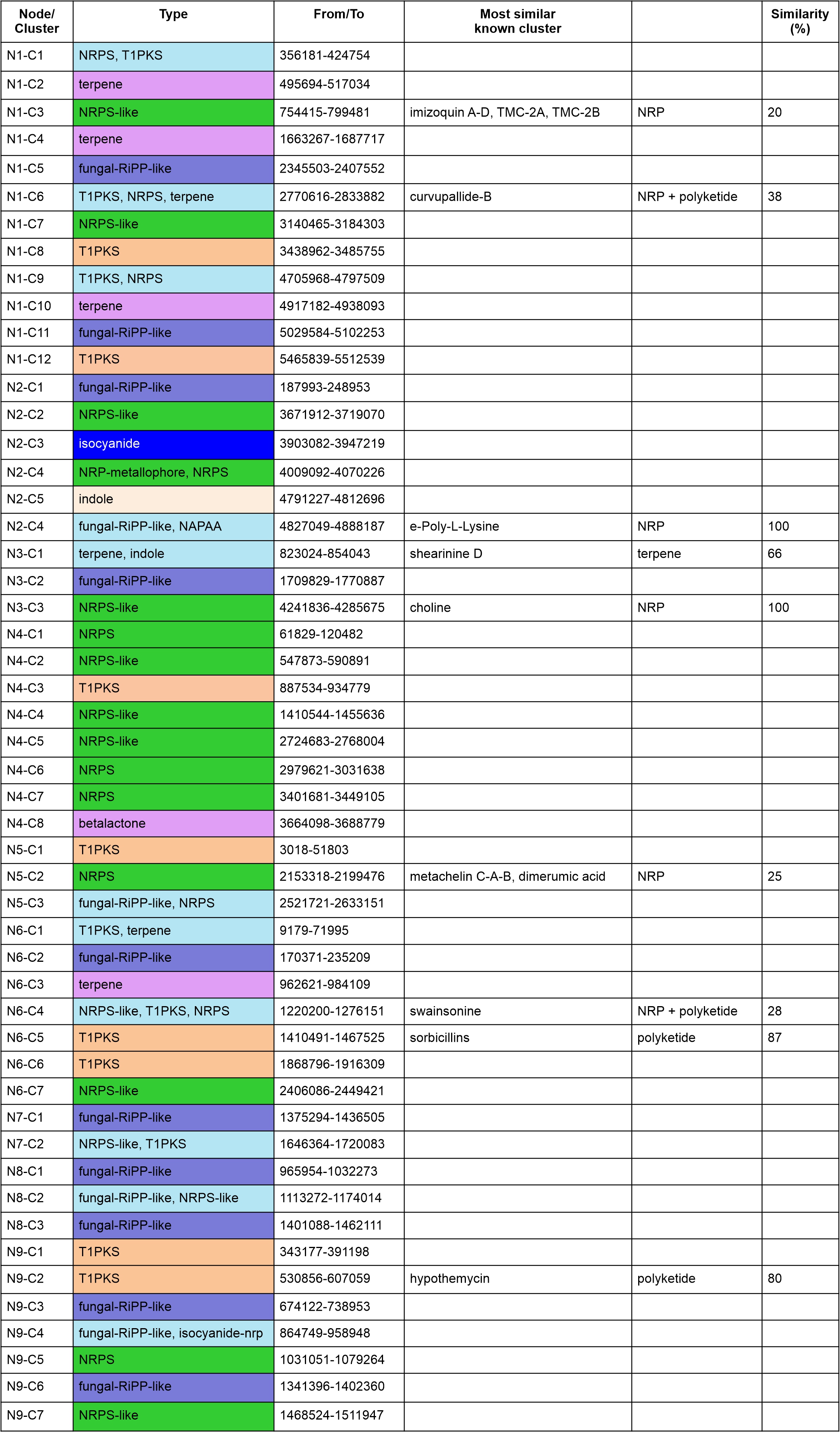

### Figure S7B

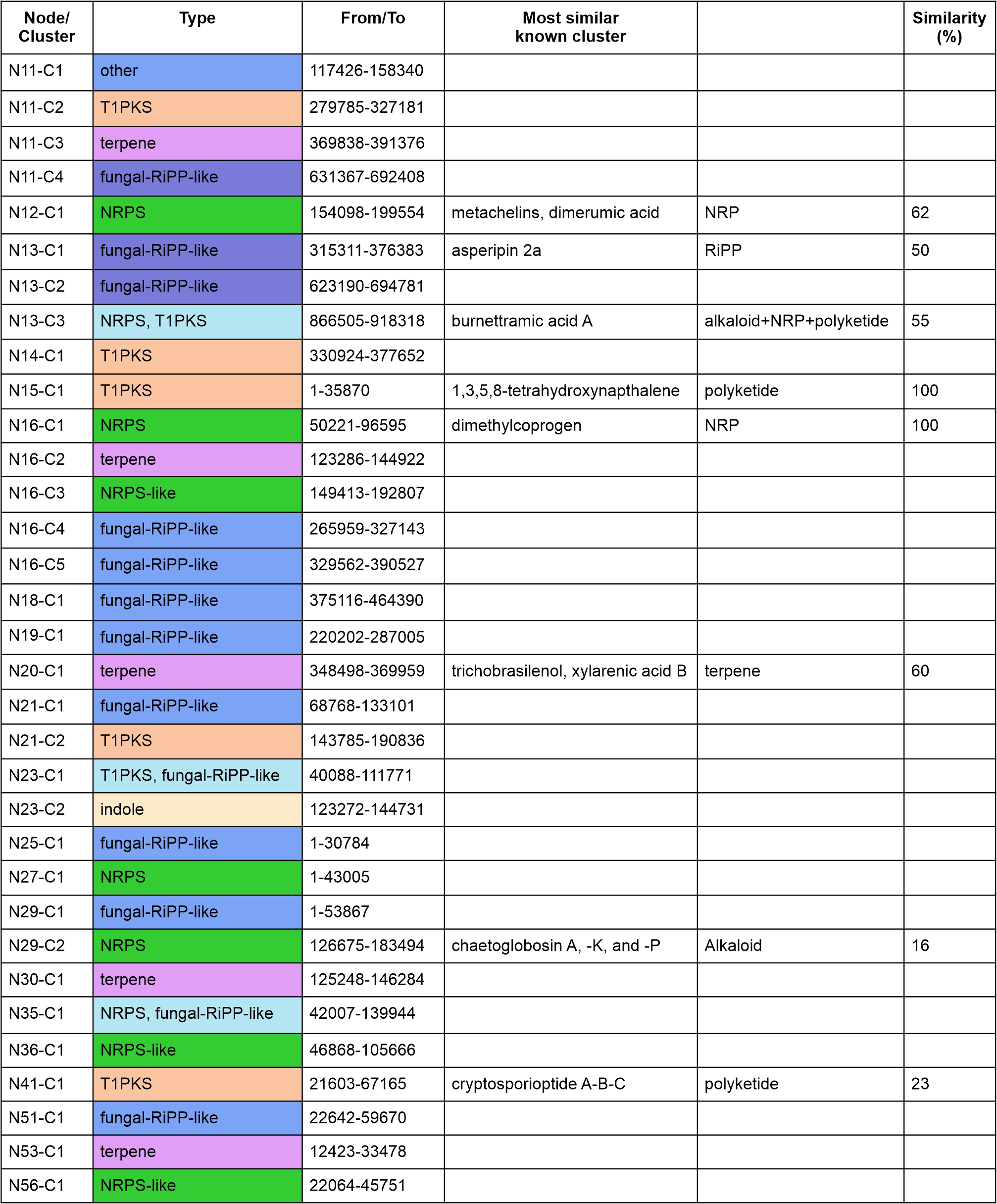

### Figure S8

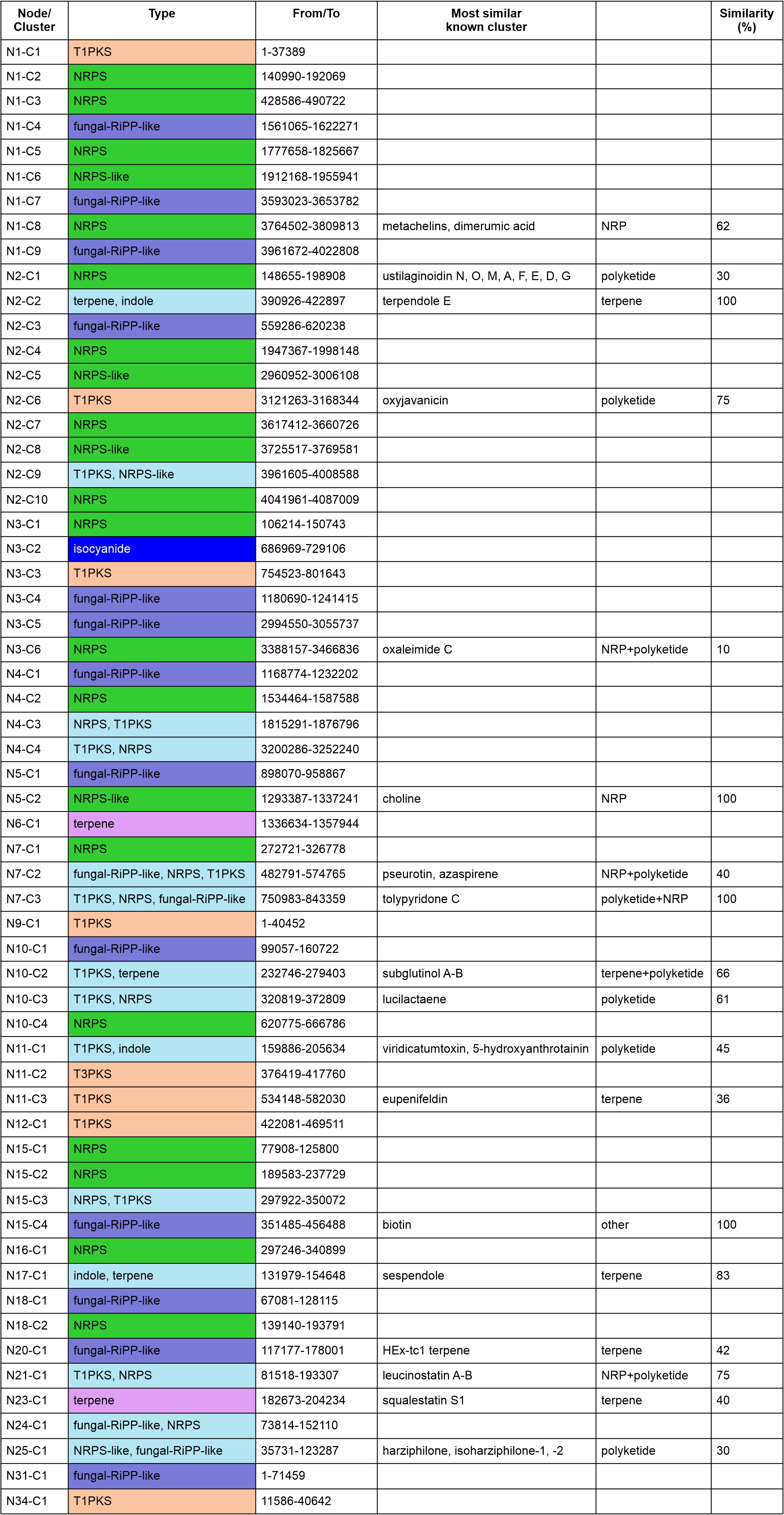

### Figure S9

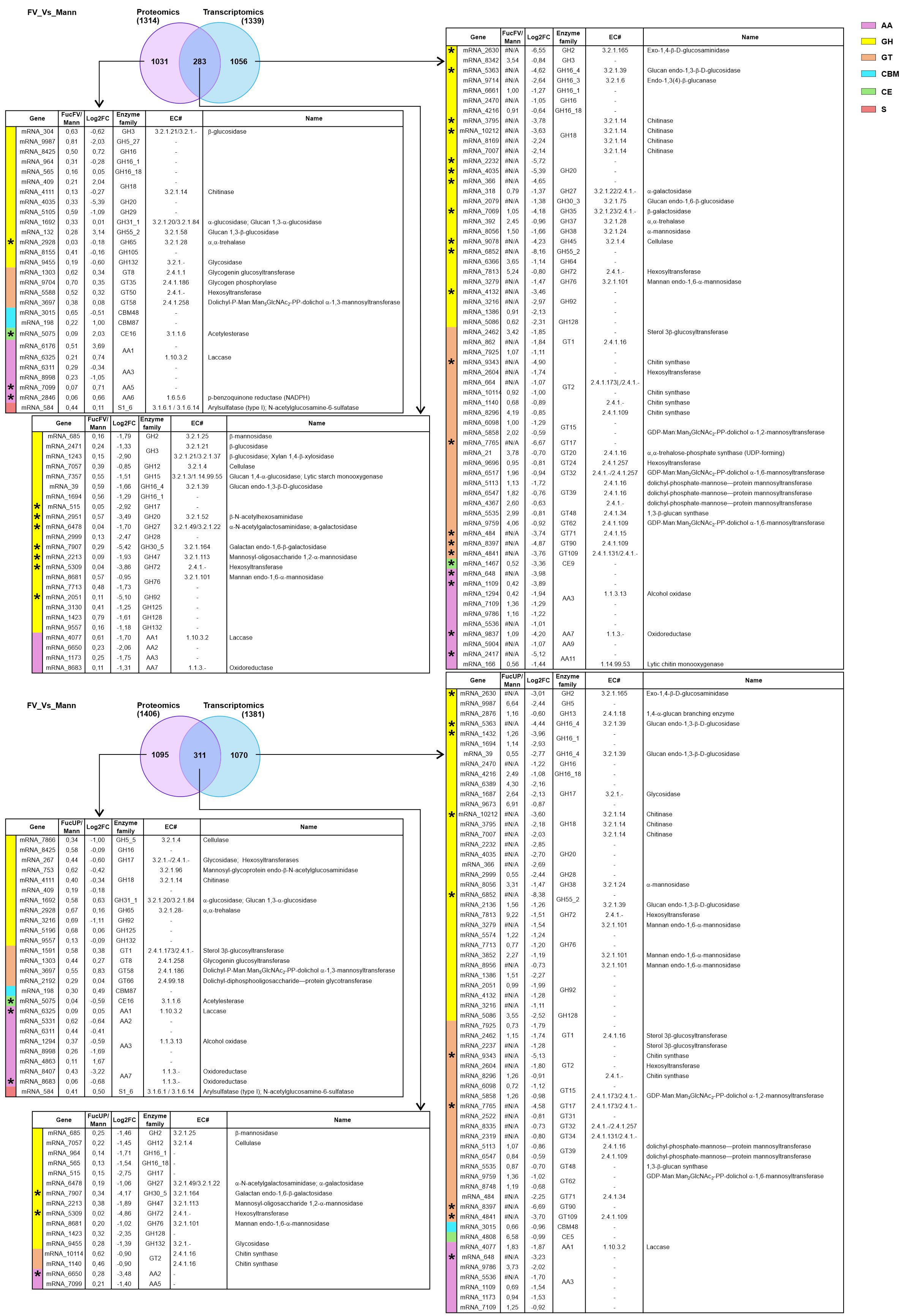
