## Supplementary material for "Phenotypic and *omics* analyses of the Sordariomycetes *Marquandomyces marquandii* and *Albophoma yamanashiensis* isolated from estuarine sediments": Table S1

| **Hypocreales Species/strain** | **GenBank Acc. Num.** | **BioSample Acc. Num.** | **Ref.** |
| --- | --- | --- | --- |
| *Acremonium chrysogenum* ATCC11550 | JPKY01000001.1 | SAMN02799700 | [1] |
| *Albophoma yamanashiensis* M98 | JAUIRT000000000 | SAMN36274349 | This study |
| *Albophoma yamanashiensis* JCM 11844 | BCKH01000001.1 | SAMD00028762 | [2], [3] |
| *Beauveria bassiana* ARSEF 8028 | JRHA01000001.1 | SAMN03066967 | [4] |
| *Calonectria pseudoturangicola* CMW 47496 | VTGA01000100.1 | SAMN12647422 | [5] |
| *Claviceps purpurea* LM5 | SRRE01000001.1 | SAMN11159852 | [6,7] |
| *Drechmeria coniospora* ARSEF 6962 | CM004174.1 | SAMN03387934 | [8] |
| *Fusarium beomiforme* NRRL 25174 | PVQB02000001.1 | SAMN07943531 | [9] |
| *Fusarium fujikuroi* NRRL 66331 | JABSTJ010000001.1 | SAMN14069559 | [9] |
| *Fusarium graminearum* NRRL28336 | LHUD01000214.1 | SAMN03958818 | [10] |
| *Fusarium oxysporum* NRRL 39464 | JAAFOW010000001.1 | SAMN13683627 | [9] |
| *Hirsutella minnesotensis* 3608 | KQ030498.1 | SAMN02777955 | [11] |
| *Hirsutella thompsonii* MTCC6686 | APKU01000001.1 | SAMN02981516 | [12] |
| *Hirsutella rhossiliensis* OWVT-1 | MPJM01000001.1 | SAMN05980824 | [13] |
| *Marquandomyces marquandii* M60 | JAUIRS000000000 | SAMN36274215 | This study |
| *Metarhizium acridum* ARSEF 324 | JAGRQL010000009.1 | SAMN18235592 | [14] |
| *Metarhizium anisopliae* ARSEF 549 | AZNF01000001.1 | SAMN03268434 | [15] |
| *Metarhizium robertsii* ARSEF 2575 | JELW01000001.1 | SAMN02798139 | [16] |
| *Neonectria coccinea* NRRL 20485 | JABSTC010000001.1 | SAMN14901119 | [9] |
| *Neonectria ditissima* R09/05 | LKCW01000001.1 | SAMN03975979 | [17] |
| *Ophiocordyceps australis* CCMB661 | JACJUF010000971.1 | SAMN15484363 | [18] |
| *Ophiocordyceps camponoti-floridani* EC05 | JAACLJ010000001.1 | SAMN13721827 | [19] |
| *Ophiocordyceps camponoti-leonardi* BCC 80369 | PDHP01000100.1 | SAMN07662903 | [20] |
| *Ophiocordyceps camponoti-rufipedis* Map16 | NJES01000001.1 | SAMN07142922 | [21] |
| *Ophiocordyceps sinensis_1* IOZ07 | JAAVMX010000001.1 | SAMN14168421 | [22] |
| *Ophiocordyceps sinensis_2* ZJB12195 | LWBQ01000001.1 | SAMN04550853 | [23] |
| *Ophiocordyceps unilateralis* SC16a | LAZP02000001.1 | SAMN03465111 | [24] |
| *Pochonia chlamydosporia_1* 170 | CM008055.1 | SAMN02296946 | [25] |
| *Pochonia chlamydosporia_2* 123 | AOSW04000001.1 | SAMN02981506 | [26] |
| *Polycephalomyces sp.* Field(B)_6/19/19 | JADHZA010000001.1 | SAMN16516720 | [27] |
| *Purpureocillium lilacinum_1* IFM 63780 | BQKX01000001.1 | SAMD00236477 | [28] |
| *Purpureocillium lilacinum_2* TERIBC 1 | LOFA01000001.1 | SAMN03701337 | [29] |
| *Purpureocillium takamizusanense* PT3 | CP086354.1 | SAMN22138285 | [30] |
| *Rugulonectria rugulosa* CBS 126565 | JALNIO010000001.1 | SAMN27671418 | [31] |
| *Stachybotrys chartarum* IBT 40293 | KL650165.1 | SAMN01819008 | [32] |
| *Tolypocladium álbum* CBS 869.73 | JALHCH010000341.1 | SAMN26764057 | [33] |
| *Tolypocladium amazonense* MS503 | JALHCD010000099.1 | SAMN26764061 | [33] |
| *Tolypocladium capitatum* CBS 113982 | NRSZ01000551.1 | SAMN07514617 | [34] |
| *Tolypocladium cylindrosporium* CBS 718.70 | JALHCC010000689.1 | SAMN26764062 | [33] |
| *Tolypocladium inflatum* CBS 567.84 | QEPE01000012.1 | SAMN08824660 | [35] |
| *Tolypocladium guangdongense* GD1-15 | NRQP01000001.1 | SAMN07540323 | [36] |
| *Tolypocladium ophioglossoides* CBS 100239 | LFRF01000001.1 | SAMN03782320 | [37] |
| *Tolypocladium ovalisporum* CBS 700.92 | JALHBZ010000069.1 | SAMN26764065 | [33] |
| *Tolypocladium paradoxum* NRBC 100945 | PKSG01000596.1 | SAMN08279508 | [34] |
| *Trichoderma reesei* QM6a | GL985056.1 | SAMN02746107 | [38] |
| *Trichoderma harzianum* CBS 226.95 | KZ679675.1 | SAMN00761861 | [39] |
| *Trichoderma cornu-damae* KA19-0412C | CM036087.1 | SAMN19791815 | [40] |
| *Ustilaginoidea virens* IPU010 | BBTG02000001.1 | SAMD00024747 | [41] |

Figure 5A: All these genomes were retrieved from <https://www.ncbi.nlm.nih.gov/>.

| **Species/strain** | **Link to MycoCosm** | **Ref.** |
| --- | --- | --- |
| *Albophoma yamanashiensis* M98 | https://mycocosm.jgi.doe.gov/Albya1/Albya1.home.html | This study |
| *Claviceps purpurea* C20.1 | https://mycocosm.jgi.doe.gov/Clapu1/Clapu1.home.html | [42] |
| *Drechmeria coniospora* ARSEF 6962 | https://mycocosm.jgi.doe.gov/Dreco1/Dreco1.home.html | [8] |
| *Hirsutella minnesotensis* 3608 | https://mycocosm.jgi.doe.gov/Hirmi1/Hirmi1.home.html | [11] |
| *Marquandomyces marquandii* M60 | https://mycocosm.jgi.doe.gov/Marma1/Marma1.home.html | This study |
| *Metarhizium album* ARSEF 1941 | https://mycocosm.jgi.doe.gov/Metal1/Metal1.home.html | [15] |
| *Metarhizium anisopliae* ARSEF 549 | https://mycocosm.jgi.doe.gov/Metani1/Metani1.home.html | [15] |
| *Metarhizium robertsii ARSEF 23* | https://mycocosm.jgi.doe.gov/Metro1/Metro1.home.html | [15] |
| *Moelleriella libera* RCEF2490 | https://mycocosm.jgi.doe.gov/Moeli1/Moeli1.home.html | [43] |
| *Ophiocordyceps australis* Map64 | https://mycocosm.jgi.doe.gov/Ophau1/Ophau1.home.html | [21] |
| *Ophiocordyceps camponoti-rufipedis* Map16 | https://mycocosm.jgi.doe.gov/Ophca1/Ophca1.home.html | [21] |
| *Ophiocordyceps sinensis* IOZ07 | https://mycocosm.jgi.doe.gov/Ophsi1/Ophsi1.home.html | [22] |
| *Paecilomyces niveus* CO7 | https://mycocosm.jgi.doe.gov/Bysni1/Bysni1.home.html | [44] |
| *Paecilomyces variotii* CBS 101075 | https://mycocosm.jgi.doe.gov/Paevar1/Paevar1.home.html | [45] |
| *Pochonia chlamydosporia* 170 | https://mycocosm.jgi.doe.gov/Pocchl1/Pocchl1.home.html | [25] |
| *Purpureocillium lilacinum* PLFJ-1 | https://mycocosm.jgi.doe.gov/Purli1/Purli1.home.html | [25] |
| *Tolypocladium capitatum* CBS 113982 | https://mycocosm.jgi.doe.gov/Tolca1/Tolca1.home.html | [34] |
| *Tolypocladium inflatum NRRL 8044* | https://mycocosm.jgi.doe.gov/Tolinf1/Tolinf1.home.html | [46] |
| *Tolypocladium ophioglossoides* CBS 100239 | https://mycocosm.jgi.doe.gov/Tolop1/Tolop1.home.html | [37] |
| *Tolypocladium paradoxum* NRBC 100945 | https://mycocosm.jgi.doe.gov/Tolpa1/Tolpa1.home.html | [34] |
| *Ustilaginoidea virens* IPU010 | https://mycocosm.jgi.doe.gov/Ustvir1/Ustvir1.home.html | [41] |

Figure S6 and Figure 5C: KOG analyses compared data retrieved from MycoCosm (<https://mycocosm.jgi.doe.gov/mycocosm/home>).

**References.**

[38]	D. Martinez, R.M. Berka, B. Henrissat, M. Saloheimo, M. Arvas, S.E. Baker, J. Chapman, O. Chertkov, P.M. Coutinho, D. Cullen, E.G.J. Danchin, I. V Grigoriev, P. Harris, M. Jackson, C.P. Kubicek, C.S. Han, I. Ho, L.F. Larrondo, A.L. de Leon, J.K. Magnuson, S. Merino, M. Misra, B. Nelson, N. Putnam, B. Robbertse, A.A. Salamov, M. Schmoll, A. Terry, N. Thayer, A. Westerholm-Parvinen, C.L. Schoch, J. Yao, R. Barabote, M.A. Nelson, C. Detter, D. Bruce, C.R. Kuske, G. Xie, P. Richardson, D.S. Rokhsar, S.M. Lucas, E.M. Rubin, N. Dunn-Coleman, M. Ward, T.S. Brettin, Genome sequencing and analysis of the biomass-degrading fungus *Trichoderma reesei* (syn. Hypocrea jecorina), Nat. Biotechnol. 26 (2008) 553–560. https://doi.org/10.1038/nbt1403.

[39]	I.S. Druzhinina, K. Chenthamara, J. Zhang, L. Atanasova, D. Yang, Y. Miao, M.J. Rahimi, M. Grujic, F. Cai, S. Pourmehdi, K.A. Salim, C. Pretzer, A.G. Kopchinskiy, B. Henrissat, A. Kuo, H. Hundley, M. Wang, A. Aerts, A. Salamov, A. Lipzen, K. LaButti, K. Barry, I. V Grigoriev, Q. Shen, C.P. Kubicek, Massive lateral transfer of genes encoding plant cell wall-degrading enzymes to the mycoparasitic fungus *Trichoderma* from its plant-associated hosts, PLOS Genet. 14 (2018) e1007322. https://doi.org/10.1371/journal.pgen.1007322.

[40]	H.-Y. Lee, J. Jong Won, K. Young-Nam, L. Hyun, R. Hojin, S. Jwakyung, S. Yoon-Sup, K. Chang-Sun, J.-W. and Chung, The complete mitochondrial genome of the poisonous mushroom *Trichoderma cornu-damae* (Hypocreaceae), Mitochondrial DNA Part B. 7 (2022) 1899–1901. https://doi.org/10.1080/23802359.2022.2135393.

[41]	T. Kumagai, T. Ishii, G. Terai, M. Umemura, M. Machida, K. Asai, Genome sequence of *Ustilaginoidea virens* IPU010, a rice pathogenic fungus causing false smut, Genome Announc. 4 (2016) 10.1128/genomea.00306-16. https://doi.org/10.1128/genomea.00306-16.

[42]	C.L. Schardl, C.A. Young, U. Hesse, S.G. Amyotte, K. Andreeva, P.J. Calie, D.J. Fleetwood, D.C. Haws, N. Moore, B. Oeser, D.G. Panaccione, K.K. Schweri, C.R. Voisey, M.L. Farman, J.W. Jaromczyk, B.A. Roe, D.M. O’Sullivan, B. Scott, P. Tudzynski, Z. An, E.G. Arnaoudova, C.T. Bullock, N.D. Charlton, L. Chen, M. Cox, R.D. Dinkins, S. Florea, A.E. Glenn, A. Gordon, U. Güldener, D.R. Harris, W. Hollin, J. Jaromczyk, R.D. Johnson, A.K. Khan, E. Leistner, A. Leuchtmann, C. Li, J. Liu, J. Liu, M. Liu, W. Mace, C. Machado, P. Nagabhyru, J. Pan, J. Schmid, K. Sugawara, U. Steiner, J.E. Takach, E. Tanaka, J.S. Webb, E. V Wilson, J.L. Wiseman, R. Yoshida, Z. Zeng, Plant-symbiotic fungi as chemical engineers: Multi-genome analysis of the clavicipitaceae reveals dynamics of alkaloid *loci*, PLOS Genet. 9 (2013) e1003323.

[43]	Y. Shang, G. Xiao, P. Zheng, K. Cen, S. Zhan, C. Wang, Divergent and convergent evolution of fungal pathogenicity, Genome Biol. Evol. 8 (2016) 1374–1387. https://doi.org/10.1093/gbe/evw082.

[44]	M.N. Biango-Daniels, T.W. Wang, K.T. Hodge, Draft genome sequence of the patulin-producing fungus *Paecilomyces niveus* strain CO7, Genome Announc. 6 (2018) 10.1128/genomea.00556-18. https://doi.org/10.1128/genomea.00556-18.

[45]	A.S. Urquhart, S.J. Mondo, M.R. Mäkelä, J.K. Hane, A. Wiebenga, G. He, S. Mihaltcheva, J. Pangilinan, A. Lipzen, K. Barry, R.P. de Vries, I. V Grigoriev, A. Idnurm, Genomic and genetic insights into a cosmopolitan fungus, *Paecilomyces variotii* (Eurotiales), Front. Microbiol. 9 (2018). https://doi.org/10.3389/fmicb.2018.03058.

[46]	K.E. Bushley, R. Raja, P. Jaiswal, J.S. Cumbie, M. Nonogaki, A.E. Boyd, C.A. Owensby, B.J. Knaus, J. Elser, D. Miller, Y. Di, K.L. McPhail, J.W. Spatafora, The genome of *Tolypocladium inflatum*: Evolution, organization, and expression of the cyclosporin biosynthetic gene cluster, PLOS Genet. 9 (2013) e1003496. https://doi.org/10.1371/journal.pgen.1003496.
